# PI31 is an adaptor protein for proteasome transport in axons and required for synaptic development and function

**DOI:** 10.1101/364463

**Authors:** Kai Liu, Sandra Jones, Adi Minis, Jose Rodriguez, Henrik Molina, Hermann Steller

**Affiliations:** Strang Laboratory of Apoptosis and Cancer Biology, The Rockefeller University, 1230 York Avenue, New York, NY 10065, USA.; The Rockefeller University Proteomics Resource Center, The Rockefeller University, New York, NY, 10065, USA

**Keywords:** proteasomes, ubiquitin-proteasome system, protein degradation, microtubule-dependent transport, axon, synapse, neurodegenerative diseases, *Drosophila*

## Abstract

Protein degradation by the ubiquitin-proteasome system (UPS) is critical for neuronal development, plasticity and function. Neurons utilize microtubule-dependent molecular motors to allocate proteasomes to synapses, but how proteasomes are coupled to motor proteins and how this transport is regulated to meet changing demand for protein breakdown remains largely unknown. We show that the conserved proteasome-binding protein PI31 serves as an adaptor to directly couple proteasomes with dynein light chain proteins (DYNLL1/2). Inactivation of PI31 inhibits proteasome motility in axons and disrupts synaptic protein homeostasis, structure and function. Moreover, phosphorylation of PI31 at a conserved site by p38 MAP kinase promotes binding to DYNLL1/2, and a non-phosphorable PI31 mutant impairs proteasome movement in axons, suggesting a mechanism to regulate loading of proteasomes onto motor proteins. Because mutations affecting PI31 activity are associated with human neurodegenerative diseases, impairment of PI31-mediated axonal transport of proteasomes may be the root cause of these disorders.

## Introduction

Protein homeostasis (proteostasis) is essential for cellular and organismal health (Balch et al., 2008; Glickman and Ciechanover, 2002; Goldberg, 2003; Labbadia and Morimoto, 2015; Varshavsky, 2005; Vilchez et al., 2014; Wolff et al., 2014). Eukaryotic cells have two major systems for clearing unwanted, damaged and potentially toxic proteins: the ubiquitin-proteasome system (UPS) and the autophagy-lysosome system(Bento et al., 2016; Dikic and Elazar, 2018; Glickman and Ciechanover, 2002; Goldberg, 2003; Levine and Kroemer, 2008; Murata et al., 2009; Nakatogawa et al., 2009; Schmidt and Finley, 2014). The UPS carries out the degradation of the vast majority of intracellular proteins, whereas autophagy is primarily responsible for the removal of protein aggregates and damaged organelles. In the UPS, proteins are tagged for destruction by the linkage of multiple copies of ubiquitin(Finley, 2009; Hershko and Ciechanover, 1998; Varshavsky, 2012). The actual degradation of proteins in the UPS is carried out by multi-subunit protease particles termed 26S proteasomes(Baumeister et al., 1998; Collins and Goldberg, 2017; Finley, 2009; Glickman and Ciechanover, 2002). The 26S proteasomes consist of 20S core particles, which contain the active proteases, and 19S regulatory subunits(Collins and Goldberg, 2017; Murata et al., 2009).

It has been suggested that proteasomes are typically present in excess and that their capacity is not fully used, at least in the absence of stress(Asano et al., 2015). On the other hand, some cell types appear to require special mechanisms to position proteasomes to appropriate sub-cellular compartments where protein breakdown occurs. This is especially important for neurons, since they are very large, structurally complex and highly compartmentalized cells that require transport mechanisms to allocate proteasomes to sites distant from the cell body (Bingol and Schuman, 2006; Bingol and Sheng, 2011; Tai and Schuman, 2008).

Proteasome function is required for growth and pruning of both axons and dendrites during development (DiAntonio et al., 2001; Erturk et al., 2014; Kuo et al., 2005; Wan et al., 2000; Watts et al., 2003). Moreover, in mature neurons UPS-mediated protein degradation at synapses is critical for activity-dependent plasticity, learning and memory(Bingol and Sheng, 2011; Campbell and Holt, 2001; Ding et al., 2007; Hegde et al., 2014; Lee et al., 2008; Pak and Sheng, 2003; Patrick, 2006; Yi and Ehlers, 2005). Recently, proteasomes were found to insert into the plasma membrane of neurons and acutely regulate synaptic activity and calcium signaling(Ramachandran et al., 2018; Ramachandran and Margolis, 2017). Due to the considerable distances from the soma, neurons employ microtubule-based transport mechanisms to deliver proteasomes to the periphery of neurons(Gorbea et al., 2010; Hsu et al., 2015; Kreko-Pierce and Eaton, 2017; Otero et al., 2014). However, despite the critical function of proteasomes at synapses, much remains to be learned about the underlying transport mechanisms. One of the molecular motors, the dynein complex, plays a key role in motility of proteasomes in axons(Hsu et al., 2015; Kreko-Pierce and Eaton, 2017). A recent study identified dynein light chain LC8-type proteins (DYNLL1/2) as components of microtubule-dependent proteasome transport in *Drosophila* neurons (Kreko-Pierce and Eaton, 2017). However, it remains unclear how proteasomes are coupled to microtubule-based motor complexes, and how proteasome transport is regulated to meet changing demands for protein breakdown.

Here we show that the conserved proteasome-binding protein PI31 (“<u>p</u>roteasome inhibitor of <u>31</u>kD”) mediates fast axonal transport of proteasomes in *Drosophila* motor neurons by acting as an adapter between proteasomes and dynein light chain LC8-type proteins (DYNLL1/2). PI31 was originally identified based on its ability to inhibit 20S proteasome-mediated hydrolysis of peptides *in vitro* (Chu-Ping et al., 1992; McCutchen-Maloney et al., 2000; Zaiss et al., 1999). On the other hand, inactivation of PI31 impairs protein degradation during sperm maturation in *Drosophila*, and PI31 can stimulate the proteolytic capacity of cells by promoting the assembly of 19S and 20S particles into the fully active 26S proteasomes(Bader et al., 2011; Cho-Park and Steller, 2013). Similarly, the PI31 ortholog in yeast, Fub1, is required to alleviate proteotoxic stress (Yashiroda et al., 2015). However, studies in cultured HEK293 cells failed to reveal a clear effect of PI31 for proteasome activity, suggesting that PI31 function may be different in different cell types and systems(Li et al., 2014).

PI31 also binds directly and strongly to the F-box protein Ntc/FBXO7/PARK15, and this interaction is conserved from *Drosophila* to mammals(Bader et al., 2011; Kirk et al., 2008). Interestingly, mutations in *FBXO7/PARK15* impair proteasome function and cause a juvenile form of Parkinson’s Disease (PD) in humans and PD-like symptoms in mice(Conedera et al., 2016; Di Fonzo et al., 2009; Paisan-Ruiz et al., 2010; Vingill et al., 2016). Since many age-related neurodegenerative diseases, including Alzheimer’s Disease (AD) and PD, are characterized by the accumulation of protein aggregates, it is possible that mutations in *FBXO7/PARK15* impair the clearance of pathological proteins (Ballatore et al., 2007; Irvine et al., 2008; Li and Li, 2011; Oddo, 2008; Ross and Poirier, 2004; Tai and Schuman, 2008). F-box proteins serve as substrate recognition modules in multi-component E3-ligase complexes for UPS-mediated protein degradation (Bai et al., 1996; Kipreos and Pagano, 2000; Skaar et al., 2013; Skowyra et al., 1997). However, very unexpectedly for an E3 ligase, binding of FBXO7 to PI31 does not result in PI31 degradation but rather serves to protect PI31 from proteolytic cleavage (Bader et al., 2011; Vingill et al., 2016). These observations are consistent with the idea that loss of *FBXO7* leads to impaired proteasome function through inactivation of PI31, but direct evidence for this model is lacking due to insufficient knowledge about the precise function of PI31 in neurons. Moreover, PI31 was linked to AD by a genome-wide association study, and mutations in another PI31-interacting protein, Valosin-Containing Protein (VCP/P97), cause amyotrophic lateral sclerosis (ALS) (Clemen et al., 2015; Johnson et al., 2010; Sherva et al., 2011).

In order to address the function of PI31 in neurons, we investigated the consequences of PI31 inactivation in *Drosophila* motor neurons. We found a striking requirement of PI31 for synaptic structure and function. Loss of PI31 function altered the structure of presynaptic active zones and caused defects in protein homeostasis in the periphery of neurons. Significantly, we identified dynein light chain proteins (dDYNLL1/Ctp and dDYNLL2/Cdlc2) as direct binding partners of PI31, and PI31 was able to promote loading of proteasomes onto dDYNLL1/2 *in vivo* and *in vitro*.

Importantly, inactivation of PI31 blocked proteasome motility in axons. We conclude that PI31 is a direct adaptor protein to load proteasomes onto dynein complexes and thereby mediates axonal transport of proteasomes. The interaction of PI31 and DYNLL1/2 is conserved from *Drosophila* to mammals, indicating that this function is conserved. Finally, we show that phosphorylation of PI31 by the stress-activated p38 MAP kinase modulates the formation of dDYNLL1/2-proteasome complexes and proteasome movement. These findings suggest that PI31 phosphorylation functions as a molecular switch to regulate the loading of proteasome onto motors. This work also reveals a molecular link between cellular stress pathways and proteasome transport. Because mutations affecting PI31 activity are associated with human neurodegenerative diseases, impairment of PI31-mediated axonal transport of proteasomes may be the root cause of these disorders.

## Results

### Requirement of PI31 for neuronal function and synaptic structure

PI31 is a conserved proteasome-binding protein, and its expression is enriched in heads and testes in *Drosophila* (Bader et al., 2011; Chintapalli et al., 2007; Cho-Park and Steller, 2013; McCutchen-Maloney et al., 2000; Yashiroda et al., 2015). To study the role of PI31 in neurons, we used *R94G06-GAL4* to drive expression of *UAS-HA-PI31* and examined its localization in neurons. The *R94G06-GAL4* driver allows for highly specific expression in a subset of motor neurons (MN1-IB) (Figure S1A). We found that PI31 forms puncta along axons and at neuromuscular junctions (NMJs), suggesting it may have a local function there (Figure 1A).

**Figure 1.**
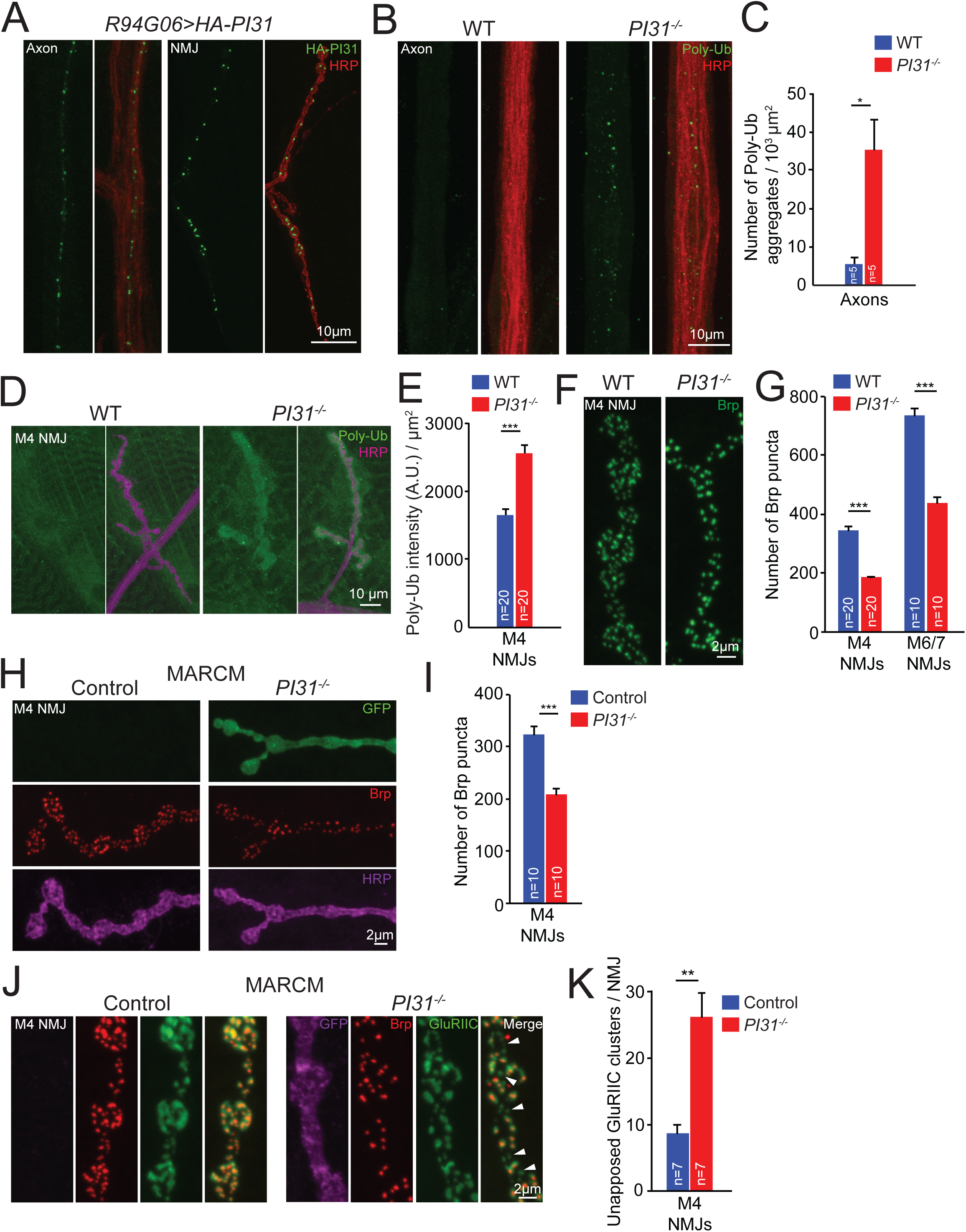
PI31 is required for proteostasis and synaptic structure and function in motor neurons. **(A)** PI31 forms puncta in axons and at neuromuscular junctions (NMJs) of *Drosophila* motor neurons. The expression of *UAS-HA-PI31* was driven by *R94G06-GAL4*, a specific motor neuron driver (Figure S1A). HA antibody (green) labels HA-PI31, and HRP antibody (red) marks neuronal membranes. **(B and C)** Representative images (**B**) and quantification(**C**) show that loss of PI31 leads to accumulation of poly-ubiquitinated (Poly-Ub) protein aggregates in axons of *Drosophila* neurons. Poly-Ub proteins were stained with a Ub-conjugates antibody (clone FK2, green), and HRP antibody (red) labeled neuronal membranes to visualize axons. Poly-Ub protein aggregates from three different microscope views were counted for each larva. The numbers were added together and normalized by the total area of axons. n=5 larvae for each genotype. Mean+s.e.m., unpaired two-tailed t-test, * *P<0.05*. **(D and E)** Representative images (**D**) and quantification (**E**) show that loss of PI31 leads to accumulation of Poly-Ub proteins at *Drosophila* NMJs. Immunofluorescence intensities of Poly-Ub proteins were measured and normalized for the NMJ area. n=20 M4 NMJs from four larvae for each genotype. Mean+s.e.m. Unpaired two-tailed t-test, *** *P<0.001*. **(F and G)** High-magnification images (**F**) and quantification (**G**) of Brp puncta at NMJs indicate that loss of PI31 leads to a structural defect of synapses. n=20 NMJs from four larvae for M4, and n=10 NMJs from six larvae for M6/7 at A2 segments. Mean+s.e.m., unpaired two-tailed t-test, *** *P<0.001*. The overview of M4 NMJs is shown in Figure S1B, and the overview and high-magnification images of M6/7 NMJs are shown in Figure S1C and S1D. **(H and I)** Representative images (**H**) and quantification (**I**) of MARCM experiments show that loss of Brp puncta at *PI31*^*-/-*^ NMJs is a cell-autonomous phenotype and indicative of a requirement of PI31 in neurons. GFP^+^ is *PI31*^*-/-*^, and GFP^-^ is *PI31*^*+/-*^ from the same larva as an internal control. n=10 M4 NMJs from eight larvae for each genotype. Mean+s.e.m., paired two-tailed t-test, *** *P<0.001*. Shown here are high-magnification images, see Figure S1F for the overview of MARCM M4 NMJs. **(J and K)** High-magnification images (**J**) and quantification (**K**) of MARCM NMJs co-stained with anti-Brp (red) and anti-GluRIIC (green, a marker for postsynaptic densities) antibodies show that some synapses at *PI31***^*-/-*^** NMJs lost Brp puncta but maintained postsynaptic densities (white arrowheads). The GFP-positive neuron (shown in a pseudo magenta color) is *PI31*^*-/-*^, whereas the GFP-negative neuron is *PI31*^*+/-*^. n=7 M4 NMJs from seven larvae for each genotype. Mean+s.e.m., paired two-tailed t-test, ** *P<0.01*. See also Figure S1.

Local activity of proteasomes plays a key role in turnover of synaptic proteins and regulates the growth, strength, activity and plasticity of synapses (Bingol and Schuman, 2006; DiAntonio et al., 2001; Ding et al., 2007; Djakovic et al., 2009; Ehlers, 2003; Erturk et al., 2014; Hegde et al., 2014; Kuo et al., 2005; Speese et al., 2003; Willeumier et al., 2006). To investigate a possible role of PI31 for protein degradation in axons and synapses, we conducted immunofluorescence studies to compare the amounts of poly-ubiquitinated proteins (Poly-Ub), the substrates of proteasomes, in axons and at NMJs of wild-type and *PI31*^*-/-*^ larvae. Quantification of these results demonstrates that *PI31*^*-/-*^ larval axons contained significantly more Poly-Ub aggregates compared to wild-type axons (Figure 1B and 1C). Likewise, the intensity of Poly-Ub staining at NMJs was also elevated (Figure 1D and 1E). This indicates that PI31 has a critical role for protein degradation at nerve terminals.

Since local activity of the UPS is important for the development of NMJs, we examined NMJ anatomy of wild-type and *PI31*^*-/-*^ larvae(DiAntonio et al., 2001; Valakh et al., 2012; Wan et al., 2000). For this purpose, we conducted immunofluorescence analyses using anti-Horseradish peroxidase (HRP) and anti-Bruchpilot (Brp) antibodies, and analyzed Muscle 4 (M4) and Muscle 6/7 (M6/7) NMJs. The anti-HRP antibody binds to neuronal membranes and serves as a marker for neuronal morphology(Jan and Jan, 1982). Brp is a core scaffold protein for presynaptic active zone (AZ) and has been widely used as a marker for proper AZ assembly. It has homology to the human AZ protein ELKS/CAST/ERC and is crucial for AZ assembly, proper clustering of Ca^2+^ channels within AZs, and efficient neurotransmitter release(Kittel et al., 2006; Owald and Sigrist, 2009; Wagh et al., 2006). Although the overall morphology of NMJs was similar between wild-type and *PI31*^*-/-*^ larvae, the number of Brp puncta was significantly reduced in *PI31*^*-/-*^ larvae, suggesting a defect of AZ assembly (Figure 1F, 1G, and S1B-E). Our results indicate that PI31 is required for the proper development of synapses at NMJ.

Next we used mosaic analysis with a repressible cell marker (MARCM) to inactivate PI31 in a limited number of cells (Wu and Luo, 2006). This analysis confirms that *PI31*^*-/-*^ neurons have reduced numbers of Brp puncta and reveals a cell-autonomous requirement of PI31 in neurons for proper synaptic morphology (Figure 1H, 1I, and S1F). Interestingly, using a GluRIIC antibody that labels postsynaptic densities to co-stain MARCM NMJs with Brp antibody, we found that some synapses in *PI31*^*-/-*^ neurons lost Brp puncta but maintained postsynaptic densities (Figure 1J and 1K, indicated by white arrowheads). Since knockdown of proteasome subunits in motor neurons causes a similar phenotype, this suggests that the synaptic defects of *PI31*^*-/-*^ neurons are due to insufficient proteasome activity at synapses(Valakh et al., 2012).

We also inactivated PI31 with a neuron-specific driver, *Nrv2-GAL4*(Sun et al., 1999). *Nrv2>PI31 RNAi* homozygotes died as pharate adults that appeared overall well-developed. These mutants initiated but did not complete the eclosion process as they displayed severe leg motor defects that made it impossible for them to escape from pupal cases (Table S1). PI31 is also required for the function and long-term survival of many other neuronal cell types in *Drosophila*, including photoreceptor neurons and dopaminergic neurons (data not shown). Taken together, these results indicate that inactivation of PI31 impairs protein degradation in axons and synapses, which causes structural and functional synaptic defects.

### P31 binds to LC8-type dynein light chain proteins

To understand the mechanism by which PI31 promotes proteostasis in the periphery of neurons, we looked for new binding partners of PI31. For this purpose, we expressed N-terminal HA-tagged PI31 (HA-PI31) in flies and performed anti-HA co-immunoprecipitation (Co-IP) followed by label-free quantitative mass spectrometry (MS). Since HA-PI31 rescued the lethality of PI31 knockout flies, we conclude that the N-terminal tag did not interfere with PI31 functions(Bader et al., 2011). Consistent with the role of PI31 as a proteasome-binding protein, HA-PI31 pulled down all 20S proteasome subunits (Figure 2A and Table S2). Interestingly, two dynein light chain LC8-type proteins, Ctp (*Drosophila* homolog of dynein light chain LC8-type 1, dDYNLL1) and Cdlc2 (*Drosophila* homolog of dynein light chain LC8-type 2, dDYNLL2) were among the statistically most significant interactors (Figure 2A). Both proteins consist of 89 amino acids, share 98.9% identity, and differ in only one amino acid (Figure S2A). Since both unique peptides were identified by MS, this suggests that both proteins can bind to PI31 (Figure S2B). Dynein proteins are microtubule-dependent molecular motors that are responsible for fast intracellular transport of various cargos (Hirokawa et al., 2010; Reck-Peterson et al., 2018; Schliwa and Woehlke, 2003; Vale, 2003). Because dDYNLL1/Ctp is involved in axonal transport of proteasomes in *Drosophila* motor neurons, it is possible that PI31 mediates proteasome transport(Kreko-Pierce and Eaton, 2017).

**Figure 2.**
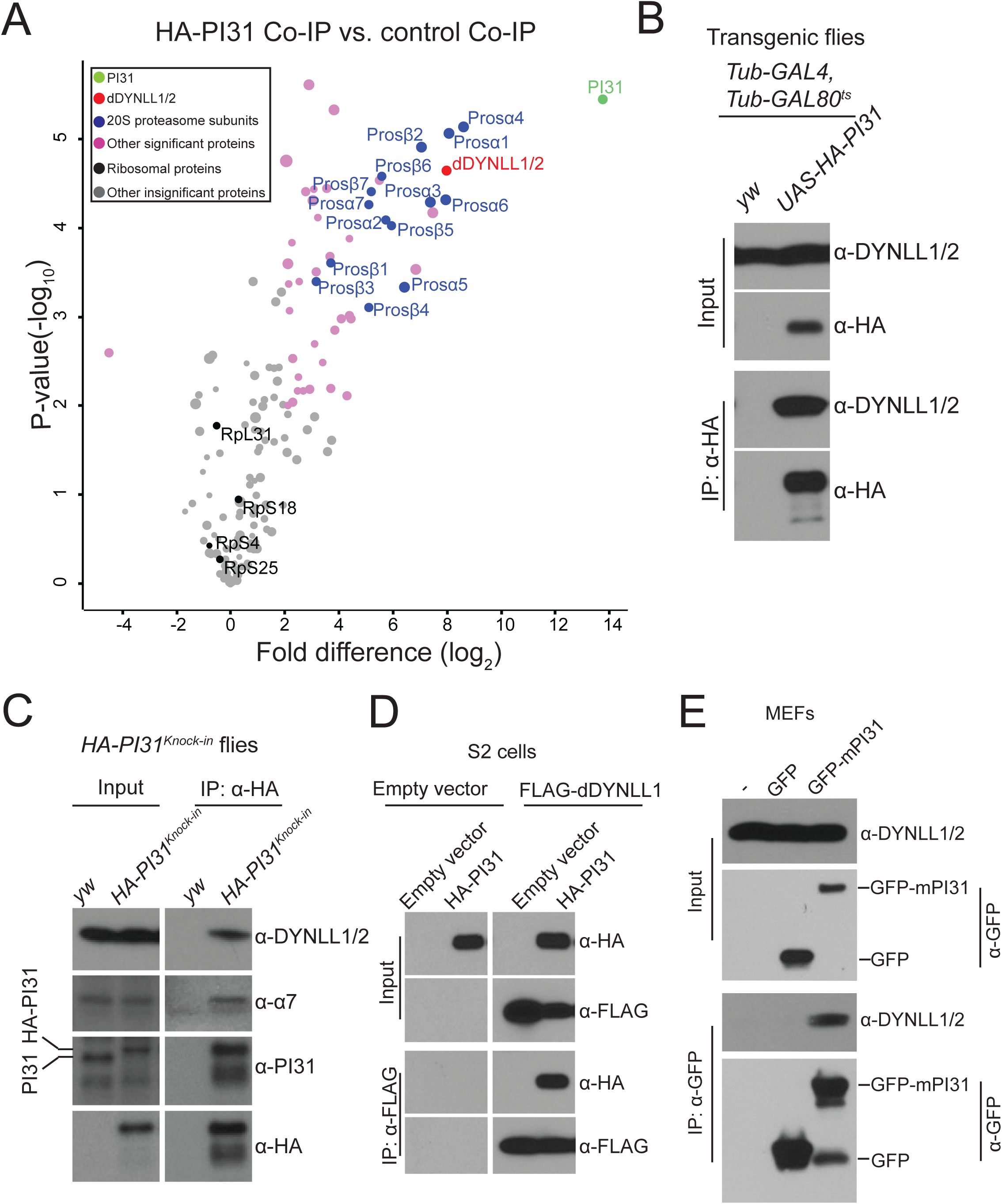
DYNLL1/2 are binding partners of PI31 in *Drosophila* and mammalian cells. **(A)** Volcano plot of quantitative MS data of anti-HA-PI31 Co-IP (*Tub-GAL4, Tub-GAL80*^*ts*^ >*UAS-HA-PI31* flies) versus control Co-IP (*Tub-GAL4, Tub-GAL80*^*ts*^ flies). Biological triplicates were conducted and analyzed. The cutoff line for significance is *P*-value<0.01(-log_10_P >2) and fold difference>4 (log_2_fold difference >2 or <-2). PI31, dDYNLL1/2, 20S proteasome subunits and other statistically significant proteins are labeled in green, red, blue and pink, respectively; whereas ribosomal proteins and other statistically insignificant proteins are labeled in black and grey, respectively. The size of dots roughly indicates the amount of a given protein detected by MS. dDYNLL1/2 are among the statistically most significant interactors of PI31, along with 20S proteasome subunits. See Table S2 for the raw data of MS analysis. **(B)** The interaction between PI31 and dDYNLL1/2 was validated by Co-IP-Western blot assays. A human DYNLL1 antibody that cross-reacts with fly dDYNLL1/2 proteins was used for Western blotting. **(C)** Co-IP-Western blot experiments using *HA-PI31*^*Knock-in*^ flies demonstrate that PI31 interacts with dDYNLL1/2 at endogenous expression levels. Anti-human DYNLL1 and proteasome subunit α7 antibodies that cross-react with homologous fly proteins were used for Western blotting. **(D)** Reciprocal Co-IP using FLAG-dDYNLL1 as the bait validates the interaction between PI31 and dDYNLL1. **(E)**The interaction between PI31 and DYNLL1/2 is conserved in mouse embryonic fibroblasts (MEFs). See also Figure S2 and Table S2.

In order to further study the interaction between PI31 and dDYNLL1/2, we conducted Co-IP-Western blot assays. HA-PI31 was able to pull down dDYNLL1/2 in both S2 cells and transgenic flies, indicating the existence of a PI31-dDYNLL1/2 complex *in vivo* (Figure 2B and S2C). To avoid possible overexpression artifacts and examine the interaction of PI31 and dDYNLL1 at endogenous expression levels, we generated a *HA-PI31*^*Knock-in*^ fly strain through CRISPR-Homology Directed Repair (HDR)(Port et al., 2014). The expression level of HA-PI31^Knock-in^ was comparable to PI31 levels in wild-type flies (Figure 2C). Using this system, we were able to show that PI31 can pull down dDYNLL1/2 and proteasomes at physiological expression levels (Figure 2C). We also performed the reciprocal Co-IP using FLAG-dDYNLL1 as the bait and found it to successfully pull down PI31 (Figure 2D).

Dynein motor complexes are highly conserved in evolution(Hirokawa et al., 2010; Reck-Peterson et al., 2018; Schliwa and Woehlke, 2003; Vale, 2003). In order to investigate whether the interaction between PI31 and dynein light chain LC8-type proteins is conserved in mammals, we performed Co-IP experiments in both MEFs and HEK293 cells and found that PI31 interacted with DYNLL1/2 in both cell types (Figure 2E and S2D). Taken together, these results indicate that PI31 interacts with DYNLL1/2 proteins, and that this interaction is conserved from *Drosophila* to mammals.

Our Co-IP-MS experiments identified 20S proteasome subunits as binding partners of PI31 but did not detect 19S proteasome subunits (Figure 2A and Table S2). However, since we did not include ATP to preserve the stability of 26S particles, this does not mean that PI31 is only in a complex with 20S proteasomes(Liu et al., 2006). To investigate whether PI31 can form a complex with 26S proteasomes, we used *HA-PI31*^*Knock-in*^ and conducted Co-IP-native gel experiments in the presence of ATP. Our results indicate that PI31 interacts with both 20S and single-capped 26S proteasomes *in vivo* (Figure S2E).

### PI31 phosphorylation stimulates direct binding to dDYNLL1/2

Co-IP-MS results revealed that *Drosophila* PI31 is phosphorylated at Serine 168 (Figure 3A and S3A). The corresponding site in human PI31 (Serine 153) is also phosphorylated, indicating that this modification is conserved from flies to human (Figure 3A and S3B). To study the function of this phosphorylation, we generated transgenic fly strains expressing a non-phosphorable mutant of *Drosophila* PI31 (Serine 168 to Alanine mutant, S168A) and compared its interactome with the interactome of wild-type PI31. Significantly, we identified dDYNLL1/2 as the most prominently affected binding partners of PI31 (Figure 3B and Table S3). This suggests that S168 phosphorylation promotes the formation of a complex between PI31 and dDYNLL1/2.

**Figure 3.**
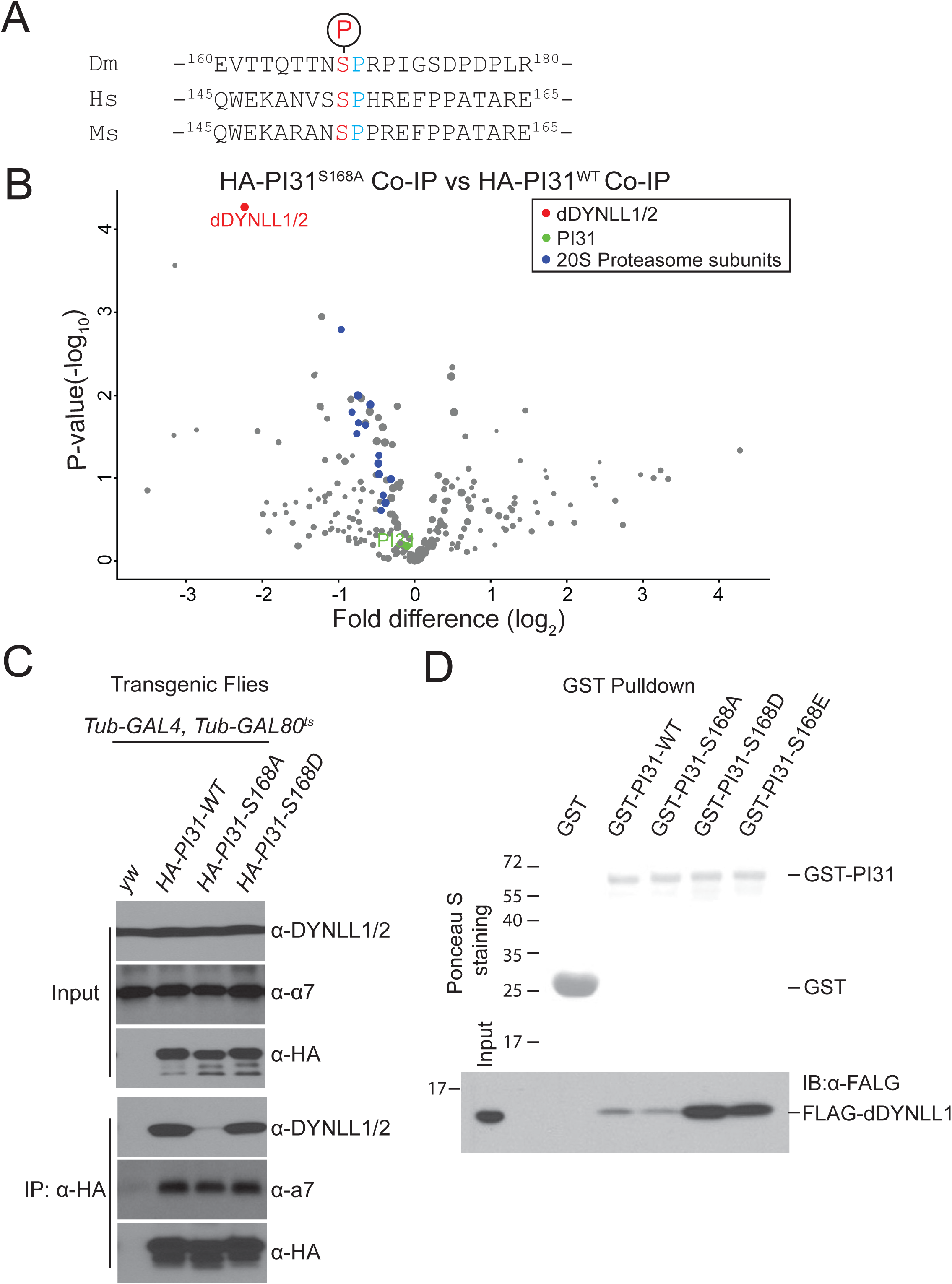
PI31 phosphorylation promotes direct interaction with dDYNLL1/2. **(A)**Alignment of the phosphorylation motifs of *Drosophila*, human and mouse PI31, indicates that PI31 is phosphorylated on a conserved serine residue. **(B)**Volcano plot of quantitative MS data of anti-HA-PI31^S168A^ Co-IP (*Tub-GAL4, Tub-GAL80*^*ts*^ >*UAS-HA-PI3* ^*S168A*^ flies) versus anti-HA-PI31^WT^ Co-IP (*Tub-GAL4, Tub-GAL80*^*ts*^ >*UAS-HA-PI3*^*WT*^ flies). Biological triplicates were conducted and analyzed. PI31, dDYNLL1/2, and 20S proteasome subunits are labeled in green, red, and blue, respectively. The size of dots roughly indicates the amount of a given protein detected by MS. The results indicate that dDYNLL1/2 are the PI31 binding partners most affected by the non-phosphorable mutation, whereas binding of 20S proteasome subunits to PI31 was not significantly changed. See Table S3 for the raw data of MS analysis. **(C)**Co-IP-Western blot analysis shows that the non-phosphorable mutation of PI31 (S168A) greatly reduces interaction with dDYNLL1/2. **(D)**GST-Pull down assays indicate that PI31 phosphorylation promotes direct interaction with dDYNLL1. All recombinant proteins were produced in *E. coli.* Ponceau S that stains total proteins was used to assess the amount and purity of recombinant GST and GST-PI31 proteins. Anti-FLAG antibody was used to detect FLAG-dDYNLL1 pulled down by different forms of GST-PI31 proteins. The results show that purified GST-PI31, but not GST, can pull down recombinant FLAG-dDYNLL1. This demonstrates that PI31 and dDYNLL1 can directly bind to each other. Furthermore, this binding was strongly enhanced by two phospho-mimetic mutations of PI31 (GST-PI31-S168D and GST-PI31-S168E), suggesting that phosphorylation of PI31 at S168 promotes complex formation. See also Figure S3 and Table S3.

To further investigate the role of PI31 phosphorylation we first conducted Co-IP-Western Blot assays. These results indicate that dDYNLL1/2 can efficiently interact with wild-type and phospho-mimetic PI31 (Serine 168 to Aspartic acid mutant, S168D) in both S2 cells and transgenic flies (Figure 3C and S3C). In contrast, this interaction was significantly reduced for the non-phosphorable mutant of PI31 (S168A) (Figure 3C and S3C). Reciprocal Co-IP using FLAG-dDYNLL1 produced similar results (Figure S3D).

Next, we investigated whether PI31 and dDYNLL1 directly interact with each other, and whether phosphorylation promotes interaction *in vitro*. For this purpose, we expressed recombinant FLAG-dDYNLL1 and GST-tagged wild-type (GST-PI31-WT), non-phosphorable mutant (GST-PI31-S168A), and two phospho-mimetic mutants (GST-PI31-S168D and GST-PI31-S168E) of PI31 proteins in *E. coli.*, and conducted GST-pull down assays. The results show that GST-PI31-WT can interact with FLAG-dDYNLL1, whereas GST alone can not (Figure 3D). Importantly, the two phospho-mimetic mutants dramatically enhanced binding, whereas the non-phosphorable mutant had no activity (Figure 3D). We conclude that PI31 is a direct binding partner of dDYNLL1, and that phosphorylation of PI31 stimulates binding.

### PI31 promotes the formation of dDYNLL1-proteasome complexes

Since PI31 can directly bind to both dDYNLL1 and proteasomes, we explored the idea that this protein may serve as an adapter to promote formation of a multi-protein complex containing proteasome and dynein. In order to look for such a complex, we expressed FLAG-dDYNLL1 in S2 cells and conducted anti-FLAG Co-IP experiments to see if proteasomes can be pulled down. Through MS analysis, we identified most of the 20S proteasome subunits (Figure 4A and Table S4). This indicates the existence of a dDYNLL1-proteasome complex. Interestingly, PI31 was among the statistically most significant binding partners of dDYNLL1, providing additional support for the relevance of this interaction. We further validated the existence of this complex by Co-IP-Western blot assays (Figure 4B).

**Figure 4.**
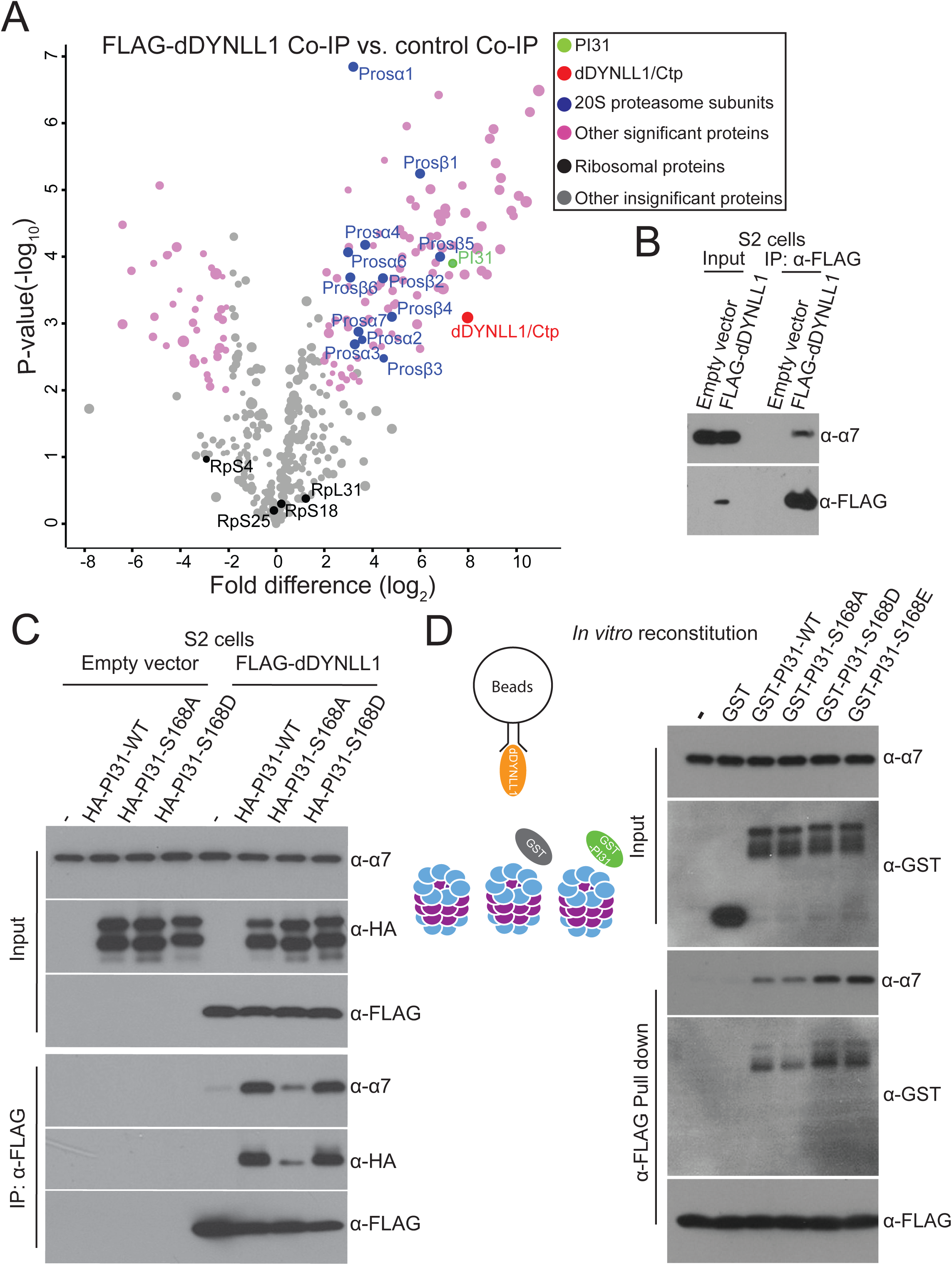
PI31 stimulates the formation of dDYNLL1-proteasome complexes. **(A)**Anti-FLAG-dDYNLL1 Co-IP and MS analysis reveals that dDYNLL1 forms a protein complex with proteasomes. S2 cells expressing FLAG-dDYNLL1 were collected for anti-FLAG Co-IP experiments. Shown here is a volcano plot of label-free quantitative MS results. Biological triplicates were conducted and analyzed. The cutoff line for significance is *P*-value<0.01(-log_10_P >2) and fold difference>4 (log_2_fold difference >2 or <-2). PI31, dDYNLL1/2, 20S proteasome subunits and other statistically significant proteins are labeled in green, red, blue and pink, respectively; whereas ribosomal proteins and other statistically insignificant proteins are labeled in black and grey, respectively. The size of dots roughly indicates the amount of a given protein detected by MS. See Table S4 for the raw data of MS analysis. **(B)** Co-IP-Western blot analysis confirms the existence of dDYNLL1-proteasome complexes. S2 cells expressing FLAG-dDYNLL1 were collected for anti-FLAG Co-IP experiments. **(C)** PI31 phosphorylation promotes the formation of dDYNLL1-proteasome complexes in S2 cells. FLAG-dDYNLL1 was expressed in S2 cells with or without different forms of HA-PI31, and then anti-FLAG Co-IP-Western blot analysis was conducted to assess proteasomes pulled down by dDYNLL1. The results indicate that PI31 can dramatically increase the amount of proteasomes bound to dDYNLL1, and this ability is largely diminished by the non-phosphorable mutation (S168A). **(D)**PI31 phosphorylation stimulates the formation of dDYNLL1-proteasome complexes *in vitro*. Recombinant FLAG-dDYNLL1 was expressed in *E. coli* and bound to anti-FLAG magnetic beads. Next, purified bovine 20S proteasomes were added alone, or with GST or different forms of GST-PI31 proteins, which were expressed and purified from *E.coli.* GST-PI31, but not GST, was able to load 20S proteasomes onto dDYNLL1 *in vitro*, and this ability was further enhanced by two phospho-mimetic mutations (S168D and S168E). The left panel depicts a schematic model for the *in vitro* binding assay, and representative results of Western Blot analyses are shown in the right panel. See also Table S4.

If PI31 is an adaptor protein mediating the formation of dDYNLL1-proteasome complexes, increased amounts of PI31 should stimulate complex formation. To test this possibility, we expressed FLAG-dDYNLL1 with different forms of HA-PI31 in S2 cells, performed anti-FLAG Co-IP experiments, and compared the amount of proteasomes pulled down by FLAG-dDYNLL1. These experiments demonstrate that elevated levels of PI31 can indeed increase the amount of proteasomes bound to dDYNLL1 (Figure 4C). Interestingly, the non-phosphorable mutant of PI31 (S168A) was not significantly active (Figure 4C).

To test whether this effect is due to direct interactions mediated by PI31, we reconstituted the complex *in vitro* using purified 20S proteasomes and recombinant GST-PI31 and FLAG-dDYNLL1 expressed in *E.coli*. First, FLAG-dDYNLL1 was bound to anti-FLAG-magnetic beads. Then, 20S proteasomes were added alone, with GST, or with GST tagged wild-type or mutant PI31 proteins. This approach showed that GST-PI31 was able to load 20S proteasomes onto dDYNLL1, whereas 20S proteasomes alone or with GST did not bind to dDYNLL1 (Figure 4D). Importantly, the two phospho-mimetic mutants (GST-PI31-S168D and GST-PI31-S168E) showed stronger ability than wild-type (GST-PI31-WT) or the non-phosphorable mutant (GST-PI31-S168A) to load proteasomes (Figure 4D). Taken together, these results show that PI31 is a direct adaptor protein that promotes the formation of a complex between proteasomes and dDYNLL1, and this activity is enhanced by phosphorylation of PI31.

### PI31 is required for the axonal transport of proteasomes in motor neurons

To examine a requirement of PI31 for axonal transport of proteasomes, we conducted live-imaging of proteasome movement in axons using the Prosβ5-RFP reporter, which is incorporated into functional proteasomes when expressed in motor neurons(Kreko-Pierce and Eaton, 2017). Expression was driven by *R94G06-GAL4* to ensure specificity for motor neurons, which is important for the determination of transport directions (Figure S1A). Inactivation of PI31 significantly reduced motility of proteasomes in axons (Figure 5A and 5B, and Movie S1 and S2). Both anterograde and retrograde movement of proteasomes was severely reduced, whereas the number of stationary proteasome particles was largely increased (Figure 5A and 5B, and Movie S1 and S2). These phenotypes are comparable to the defects described for *dDYNLL1/ctp*^*-/-*^ motor neurons, supporting the idea that they function in the same protein complex(Kreko-Pierce and Eaton, 2017). Collectively, these results demonstrate that PI31 is required for proteasome transport along motor neuron axons.

**Figure 5.**
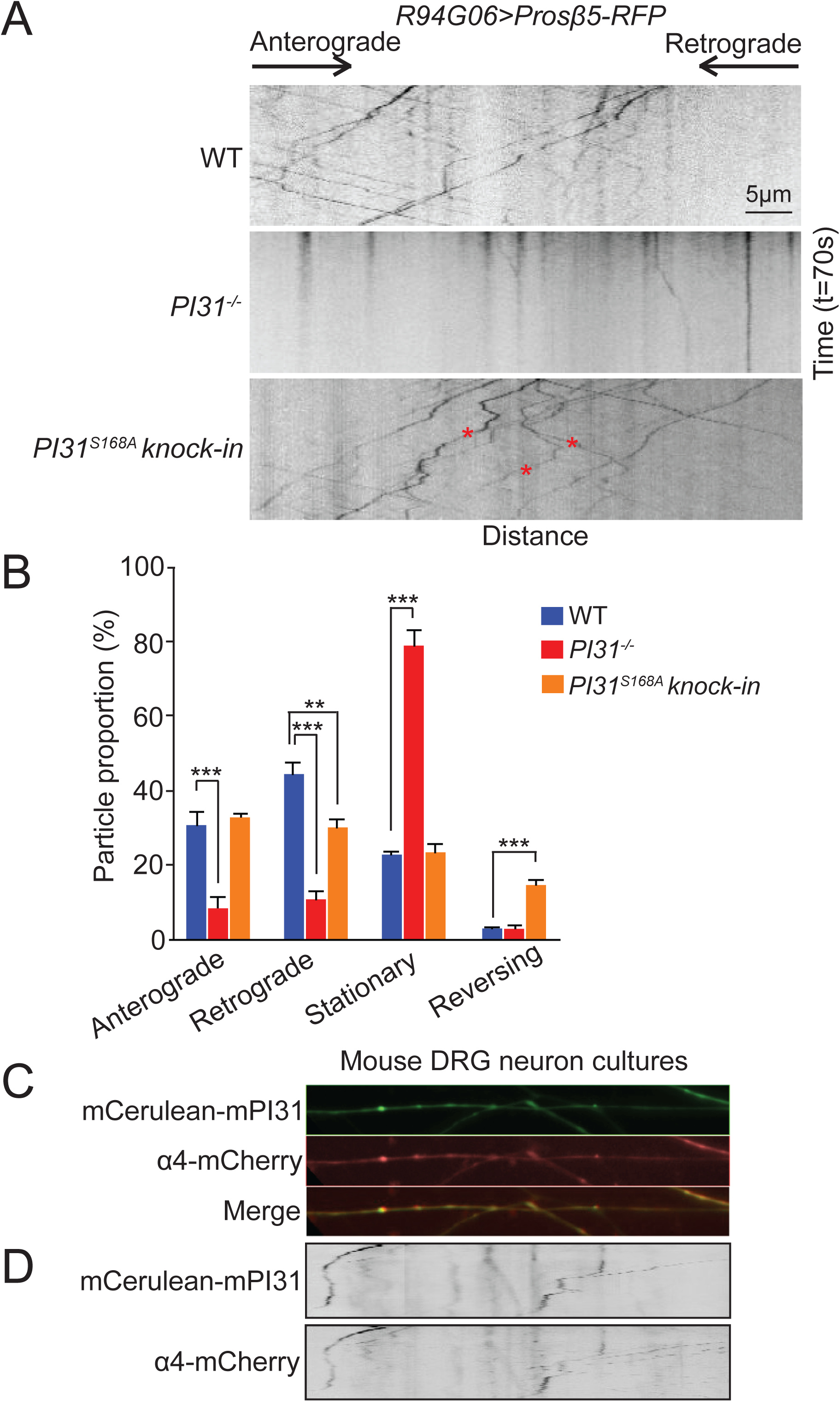
PI31 and its phosphorylation are required for axonal transport of proteasomes in motor neurons. **(A)** Representative kymographs of Prosβ5-RFP motility in motor neurons of wild-type, *PI31*^*-/-*^, and *PI31*^*S168A*^ *knock-in* larvae. *Prosβ5-RFP* expression was driven by *R94G06-GAL4*, a motor neuron-specific driver (See Figure S1). The kymographs were generated from 70-second live imaging (100 frames). The x-axis represents distance, while the y-axis represents time. The directions of anterograde and retrograde movement are indicated on top of the kymographs. Stationary particles appear as vertical lines, whereas motile particles appear as diagonal lines on kymographs. The red asterisks indicate the tracks of three proteasome particles in *PI31*^*S168A*^ *knock-in* axons that changed their moving direction during living imaging. See Figure S4 for the generation of the *PI31*^*S168A*^ *knock-in* strain. **(B)** Quantification of proteasome movement. Shown are the percentages of Prosβ5-RFP particles that moved in anterograde direction, moved in retrograde direction, appeared stationary and reversely moved in axons of wild-type, *PI31*^*-/-*^ and *PI31*^*S168A*^ *knock-in* motor neurons. Mean + s.e.m. n=7 larvae for wild-type, n=12 larvae for *PI31*^*-/-*^, and n=7 larvae for *PI31*^*S168A*^ *knock-in.* One-way Anova with Tukey’s honestly significant difference post hoc test. \*\*\**P<0.001*, ** *P<0.01*. In axons of *PI31*^*-/-*^ motor neurons, both anterograde and retrograde movements of proteasome particles were drastically reduced, while the portion of stationary proteasome particles increased accordingly; PI31^S168A^ reduced retrograde transport of proteasomes and greatly increased the frequency of proteasomes that reversed the direction of their movement. **(C and D)** Representative fluorescence images (**C**) and kymographs (**D**) of cultured mouse DRG neurons expressing mCerulean-mPI31 (green) and α4-mCherry (red) demonstrate that they extensively co-localize and move together in axons. The kymographs were generated from a 140-second live imaging experiment (70 frames). See also Figure S4 and Movies S1-4.

Our biochemical data indicate that S168 phosphorylation of PI31 is important for interaction with dDYNLL1 and enhances loading of proteasomes onto dDYNLL1. To examine the physiological role of PI31-phosphorylation for proteasome transport, we generated *PI31*^*S168A*^ *knock-in* mutants in which the wild-type PI31 gene was replaced with the non-phosphorable mutant (Figure S4). Live imaging of Prosβ5-RFP in homozygous *PI31*^*S168A*^ *knock-in* larvae revealed no significant effects on anterograde and stationary proteasome particles. However, retrograde movement of proteasome particles was significantly reduced, and we also saw a dramatic increase in the number of randomly moving particles (Figure 5A, indicated by red asterisks, Figure 5B, and Movie S3). These results indicate that PI31 phosphorylation is important for retrograde transport of proteasomes and to maintain directionality of movement.

Since PI31 interacts with proteasomes and dynein light chain LC8-type proteins in both *Drosophila* and mammals, we explored the possibility that the function of PI31 as an adapter to mediate axonal transport of proteasomes is conserved in evolution. As a first step, we examined the subcellular localization of PI31 and proteasomes in axons of cultured mouse dorsal root ganglion (DRG) neurons. This showed that α4-mCherry and mCerulean-mPI31 co-localize and move together along DRG axons (Figure 5C and 5D, and Movie S4). These observations are consistent with the idea that the function of PI31 as an adapter to mediate axonal transport of proteasomes has been conserved to mammals.

### p38 MAPK can phosphorylate PI31

In order to identify the kinase responsible for PI31 phosphorylation, we conducted a candidate-based RNAi screen. Quantitative MS results indicated that knockdown of p38β MAPK reduced PI31 phosphorylation by almost 50%, suggesting it as a candidate for PI31 phosphorylation (Figure 6A).

**Figure 6.**
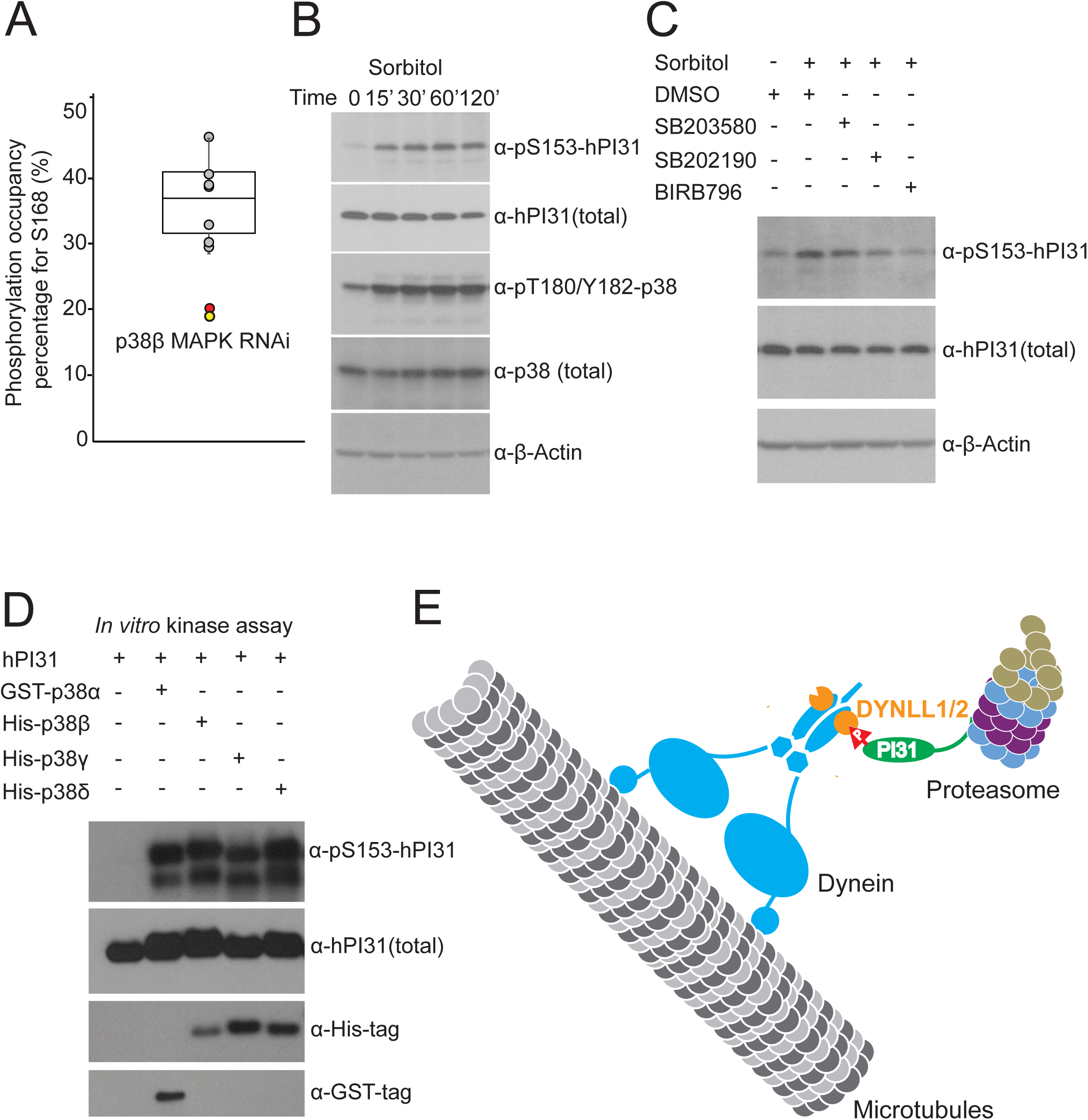
p38 MAPK can phosphorylate PI31. **(A)** A candidate-based kinase RNAi screen identified p38β as the kinase that phosphorylates PI31 at S168. *Tub-GAL4, Tub-GAL80*^*ts*^*>UAS-HA-PI31* flies were crossed with RNAi transgenic fly strains targeting each kinase of MAPK or CDK family. HA-PI31 protein was immunoprecipitated and digested. The peptides containing the phosphorylated and non-phosphorylated S168 (EVTTQTTN^168^SPRPIGSDPDPLR) were quantified by MS analysis. Phospho-peptide signal versus total signal was used to estimate the phosphorylation occupancy percentage (POP). The POP distribution of kinase RNAi strains is presented as a box plot with the following percentiles: 10% and 90% (bottom and top whiskers), 25% and 75% (bottom and top of box) and 50% (middle line). The POPs for control RNAi strains are shown as grey dots, and the POPs for replicates of the P38β RNAi are shown as red and yellow dots, suggesting ∼50% decrease of phosphorylation. See Table S5 for the summary of MS analysis. **(B)** The S153 phosphorylation of human PI31 (pS153-hPI31) increased in response to osmotic stress, a classical condition to activate p38. HEK293 cells were treated with 0.3M sorbitol for indicated time. See Figure S5 for the generation and validation of the pS153-hPI31 antibody. **(C)** The increase of pS153-hPI31 under osmotic stress was suppressed by p38 inhibitors. HEK293 cells were pretreated with indicated compounds (SB203580, SB202190 and BIRB796) at 10 µM for 2 hours, followed by the treatment of 0.3M sorbitol for 1 hour. The results show that SB203580 and SB202190, which inhibit p38α and p38β, only mildly suppressed pS153-hPI31, whereas BIRB796 that targets all four isoforms of p38, dramatically reduced pS153-hPI31. This suggests that PI31 phosphorylation requires p38 activity. **(D)** *In vitro* kinase assays using recombinant proteins generated in *E coli.* demonstrate that p38 can directly phosphorylate hPI31 at S153. **(E)** Model for PI31-mediated transport of proteasomes in axons. Proteasomes can move rapidly along microtubules to fulfill dynamic local demands for protein degradation in different cellular compartments(DiAntonio et al., 2001; Hegde et al., 2014; Hsu et al., 2015; Kreko-Pierce and Eaton, 2017; Watts et al., 2003; Zhao et al., 2003). PI31 binds directly to both proteasomes and dynein light chain LC8-type proteins (DYNLL1/2). Inactivation of PI31 severely reduces proteasome movement and leads to the accumulation of Poly-Ub aggregates in the periphery of neurons. This suggests that PI31 functions as an adaptor to directly couple proteasomes to microtubule-based motors. PI31 phosphorylation by the stress-activated p38 MAP kinase regulates the PI31-dDYNLL1 interaction and thereby modulates proteasome transport in axons. This may serve to dynamically adjust the spatio-temporal distribution of proteasomes in response to stress to match changing demands for protein breakdown. If this mechanism is impaired, mis-regulation of proteasome distribution occurs and leads to a failure of localized protein degradation. See also Figure S5 and Table S5.

p38 MAPKs can be activated by a wide range of cellular stresses as well as inflammatory cytokines(Cuenda and Rousseau, 2007; Zarubin and Han, 2005). One classical stress to activate p38 is sorbitol-induced osmotic stress(Cuenda and Rousseau, 2007; Han et al., 1994; Zarubin and Han, 2005). Therefore, we asked whether PI31 phosphorylation is elevated in response to osmotic stress. First, we generated a phosphorylation-specific antibody for Serine 153 of human PI31 (hPI31) and verified its specificity (Figure S5). Next, we treated HEK293 cells with sorbitol and assessed S153 phosphorylation (pS153) of endogenous hPI31. This showed that pS153 levels were indeed elevated after treatment, suggesting that stress promotes phosphorylation of PI31 at S153 (Figure 6B). To test whether this stress-induced phosphorylation depends on p38, we applied several distinct p38 inhibitors. Inhibition of p38α and p38β had only minor effects, but using an inhibitor targeting all four isoforms of p38 dramatically reduced S153 phosphorylation of PI31 (Figure 6C). This suggests that multiple isoforms of p38 can phosphorylate PI31. Finally, we used purified recombinant p38 and hPI31 to conduct *in vitro* kinase assays and found that all four isoforms of p38 can directly phosphorylate PI31 (Figure 6D). We conclude that p38 can phosphorylate PI31 in response to stress.

## Discussion

The degradation of unwanted and potentially toxic proteins by the ubiquitin-proteasome system (UPS) is of critical importance for protein homeostasis. In addition, UPS-mediated protein degradation also sculpts cell shape during development and modifies tissues throughout life. Examples include the terminal differentiation of spermatids, loss of muscle mass in response to starvation, dendritic pruning of nerve cells, and the maturation and modification of synapses to mediate neuronal plasticity(Bader et al., 2011; Campbell and Holt, 2001; Hegde et al., 2014; Lecker et al., 1999; Lee et al., 2008; Pak and Sheng, 2003; Patrick, 2006; Qian et al., 2013; Sandri, 2013; Watts et al., 2003; Yi and Ehlers, 2005; Zhong and Belote, 2007). Neurons pose a particular challenge for regulated protein degradation due to their large size and highly compartmentalized structure. Localized regulation of UPS activity is crucial for synaptic growth, function and plasticity, and remodeling of axons and dendrites (Bingol and Schuman, 2006; DiAntonio et al., 2001; Djakovic et al., 2012; Ehlers, 2003; Erturk et al., 2014; Hamilton and Zito, 2013; Kuo et al., 2005; Pak and Sheng, 2003; Ramachandran and Margolis, 2017; Speese et al., 2003; Wan et al., 2000; Watts et al., 2003; Zhao et al., 2003). Nerve cells employ microtubule-based transport mechanisms to quickly distribute proteasomes between soma and neurites(Bingol and Schuman, 2006; Hsu et al., 2015; Kreko-Pierce and Eaton, 2017; Otero et al., 2014). Recent reports suggested that dynein complexes, especially the dynein light chain LC8-type proteins (DYNLL1/2), are involved in microtubule-dependent proteasome transport in neurons(Hsu et al., 2015; Kreko-Pierce and Eaton, 2017). However, central unresolved questions are how proteasomes are coupled to microtubule-based motor complexes, and how this process is regulated to meet dynamic demands of localized protein degradation. Advances in this area are important since they provide the foundation for specific manipulation of proteasome transport, including to potentially boost clearance of toxic proteins in the periphery of neurons.

One candidate protein as a transport adaptor for proteasomes is Ecm29(Hsu et al., 2015). This study used a fluorescent reporter, MV151, to visualize proteasome movement in axons of cultured hippocampal neurons. One concern with the use of this compound is that MV151 binds to catalytic β-subunits and inhibits proteasome activity(Verdoes et al., 2006). Significantly, Ecm29 is a proteasome quality control factor that is recruited to aberrant proteasomes (De La Mota-Peynado et al., 2013; Finley et al., 2016; Lee et al., 2011; Leggett et al., 2002; Lehmann et al., 2010; Park et al., 2011; Wang et al., 2010). Therefore, an alternative interpretation is that Ecm29-modulated proteasome movement may represent a quality control mechanism in response to proteasome inhibition by MV-151. Moreover, for Ecm29 to qualify as an “adapter” it remains to be convincingly demonstrated that Ecm29 can bind directly to motor proteins. Therefore, the precise role of Ecm29 in the formation of proteasome-motor complexes and the physiological function of this protein for neuronal development and function *in vivo* deserves further examination.

Here we identified dynein light chain LC8-type proteins (DYNLL1/2) as direct interactors of the conserved proteasome-binding protein PI31 (“<u>P</u>roteasome <u>I</u>nhibitor of <u>31</u>kD”). This raised the possibility that PI31 serves as an adapter to couple proteasomes via dynein light chain proteins to microtubule-based motors. In support of this idea, PI31 promoted the formation of proteasome-DYNLL1/2 complexes *in vivo* and *in vitro*, and inactivation of PI31 disrupted axonal proteasome movement and caused functional and structural defects of synapses in *Drosophila*. Furthermore, loss of PI31 function led to the accumulation of Poly-Ub aggregates at nerve terminals and in axons, consistent with the idea that insufficient amounts of proteasomes were available in the periphery of neurons for proper protein degradation. Collectively, our results suggest that PI31 is a direct adaptor for microtubule-dependent transport of proteasomes and thereby regulates allocation of proteasomes between soma and synapses (Figure 6E).

Our work also revealed a role of p38 MAPK-mediated phosphorylation of PI31 in the formation of proteasome-DYNLL1/2 complexes. Phosphorylation of a conserved site in PI31 promoted binding to DYNLL1/2 and stimulated the formation of proteasome-DYNLL1/2 complexes. Importantly, mutating the phospho-acceptor site of PI31 (*PI31*^*S168A*^ *knock-in*) impaired proteasome transport in axons. This suggests that PI31 phosphorylation serves as a molecular switch to regulate proteasome movement (Figure 6E). Since phosphorylation is a dynamic and labile post-translational modification, PI31 may use phosphorylation to receive various internal and external signals. p38 MAPKs can be activated by a wide range of cellular stresses, including proteotoxic stress(Cuenda and Rousseau, 2007; Zarubin and Han, 2005). Our findings are consistent with the idea that p38 MAPKs stimulate proteasome movement in response to stress by promoting the formation of proteasome-DYNLL1/2 complexes. Although both *PI31*^*-/-*^ and *PI31*^*S168A*^ *knock-in* mutants showed impairment of proteasome transport in axons, their phenotypes were not identical. Whereas complete inactivation of PI31 led to striking impairment of both anterograde and retrograde proteasome movement, the major defects in the *PI31*^*S168A*^ mutant were reduced retrograde movement and loss of directionality (i.e., the number of proteasomes changing direction was strikingly increased compared to wild type). Presumably the most likely explanation for this difference is that *PI31*^*S168A*^ is a hypomorphic mutant, which reduces but does not completely abolish the interaction between PI31 and dDYNLL1/2. However, we cannot rule out that PI31 interacts with other motor proteins through phosphorylation-independent mechanisms, and/or that other post-translational modifications affect this process.

This study may help reconcile somewhat conflicting reports in the literature. PI31 was initially identified as a proteasome-interacting protein, and binding of PI31 to 20S particles can inhibit peptide proteolysis *in vitro*(Bader et al., 2011; Chu-Ping et al., 1992; McCutchen-Maloney et al., 2000; Zaiss et al., 1999). On the other hand, genetic studies in both yeast and *Drosophila* indicate that the physiological function of PI31 is to promote protein breakdown(Bader et al., 2011; Yashiroda et al., 2015). It is also worth noting that PI31 activity is subject to post-translational modification, and that large cells, such as spermatids and neurons are particular sensitive to the loss of PI31(Bader et al., 2011; Cho-Park and Steller, 2013). In contrast, no overt requirements for PI31 were detected in cultured HEK293 cells, and PI31-inactivation produces only modest biochemical changes at the level of whole-cell extracts(Cho-Park and Steller, 2013; Li et al., 2014). The mechanism reported here provides an explanation for these seemingly conflicting reports as it reveals a role of PI31 for localized regulation of proteasomes. Such a localized function is not readily detectable by standard methodologies for assessing proteasome activity in whole-cell extracts.

The results presented here may also have important implications for human neurological disorders. Many age-related neurodegenerative diseases, including Alzheimer’s Disease (AD) and Parkinson’s Disease (PD), are characterized by the accumulation of protein aggregates(Ballatore et al., 2007; Irvine et al., 2008; Li and Li, 2011; Oddo, 2008; Ross and Poirier, 2004; Tai and Schuman, 2008). The proteins in these pathognomonic aggregates are typically ubiquitin-conjugated (Bilen and Bonini, 2005; de Vrij et al., 2004; Mori et al., 1987; Morimoto, 2008; Morishima-Kawashima et al., 1993; Perry et al., 1987; Rubinsztein, 2006; Zoghbi and Orr, 2000). This suggests that the affected neurons attempted to remove abnormal proteins by tagging them with ubiquitin, but subsequently failed to degrade them. One possible explanation for this is that proteasomes were not available in sufficient numbers to degrade all ubiquitin-conjugates. Because the first signs of impaired proteostasis and accumulation of pathological proteins are generally seen in the periphery of neurons, it has been suggested that defects in proteasome transport may be responsible for abnormal protein clearance in neurodegenerative diseases(Coleman and Perry, 2002; Gendron and Petrucelli, 2009; Hoover et al., 2010; Ittner et al., 2010; Otero et al., 2014; Yoshiyama et al., 2007; Yu and Lu, 2012). PI31 is implicated in this process since human mutations affecting PI31 activity are associated with neurodegenerative diseases, including AD, PD and amyotrophic lateral sclerosis (ALS)(Conedera et al., 2016; Di Fonzo et al., 2009; Johnson et al., 2010; Koppers et al., 2012; Paisan-Ruiz et al., 2010; Sherva et al., 2011). Perhaps the most compelling case for linking PI31 with human disease comes from the analysis of *FBXO7/PARK15* mutants. Mutations in this gene lead to an early, familiar form of PD in humans, and inactivation of FBXO7 in mice causes neuronal degeneration and PD-like phenotypes (Conedera et al., 2016; Di Fonzo et al., 2009; Paisan-Ruiz et al., 2010; Vingill et al., 2016). FBXO7 is a strong and direct binding partner of PI31 in both insects and mammals, and loss of FBXO7 function results in proteolytic cleavage and reduced levels of PI31(Bader et al., 2011; Kirk et al., 2008). Strikingly, *FBXO7*-null mice were found to have significantly reduced PI31 levels, but a critical function of PI31 was dismissed based on analysis of whole-cell extracts(Vingill et al., 2016). Since a role of PI31 in proteasome transport would not have been detected by this approach, we suggest that this topic deserves further investigation. Finally, p38 MAPKs are involved in synaptic plasticity, learning and memory(Giese and Mizuno, 2013). They have also been implicated in the pathogenesis of various human neurological diseases, including AD, PD, ALS, ischemia, neuropathic pain, epilepsy and depression (Cuenda and Rousseau, 2007; Jha et al., 2015; Kim and Choi, 2010; Lee and Kim, 2017; Munoz and Ammit, 2010; Yasuda et al., 2011). Our results link p38 MAPKs with the regulation of proteasome transport and raise the intriguing possibility that mis-regulation of proteasome transport contributes to the pathogenesis of p38 MAPK-associated neurological disorders.

## Acknowledgement

We thank Dr. Aaron DiAntonio for the GluRIIC antibody; Dr. Benjamin Eaton, the Bloomington *Drosophila* Stock Center, Vienna *Drosophila* Resource Center, FlyLight team of Janelia Research Campus, and Flybase for *Drosophila* strains and relevant information. We are also grateful to Drs. Park Cho-Park, Simon Bullock, Michael Davidson, Didier Trono and Steven Vogel, and Addgene for plasmids. We thank Yetis Gultekin for generating the pLVX-EF1a-AcGFP-C1-mPI31 plasmid and anti-PI31 antibody, and Drs. Alison North, Christina Pyrgaki and Tao Tong for their help on live-imaging analysis. We also thank Drs. Michael Young, Brian Chait, Sarit Larisch, Park Cho-Park, Xiaochun Li, Mia Horowitz, Avi Levin, Sigi Benjamin-Hong, Dolors Ferres-Marco, Junko Shimazu, Thomas Hsiao, Wanhe Li, Erica Jacobs, and Alana Persaud for valuable discussions. This work was supported by NIH grant RO1GM60124 and a generous gift from the Loewenberg Foundation to H.S.

## Author Contributions

Conceptualization, K.L. and H.S.; Investigation, K.L., S.J., A.M., J.R., and H.M.; Resources, H.S.; Writing, K.L. and H.S.; Funding Acquisition, H.S.; Supervision, H.S.

## Methods

### Fly husbandry and strains

All flies were raised on standard molasses formulation food at 25°C with a 12-hour light/12-hour dark cycle. The following fly strains were used: *R94G06-GAL4* (Janelia GAL4 collection, Bloomington Drosophila Stock Center, BDSC#40701) (Pfeiffer et al., 2008), *UAS-mCD8-mCherry* (BDSC #27392), *UAS-HA-PI31* (2^nd^ chromosome)(Bader et al., 2011), *PI31*^*-/-*^(Bader et al., 2011), *hsp70-FLP*^*1*^, *FRT42D tubP-GAL80* (BDSC #9917), *D42-GAL4* (BDSC # 8816), *UAS-IVS-myr::GFP* (BDSC #32197) (Pfeiffer et al., 2010), *tubP-GAL4* (BDSC # 5138), *tubP-GAL80*^*ts*^ (BDSC #7018), *Nrv2-GAL4 UAS-GFP* (BDSC #6794), *UAS-PI31 RNAi*^*KK105476*^ (Vienna Drosophila Resource Center, #105476) and *UAS-Prosβ5-RFP*(Kreko-Pierce and Eaton, 2017) (kindly provided by Dr. Benjamin A. Eaton, University of Texas). *UAS-HA-PI31-WT* ^*attP_ ZH-68E*^, *UAS-HA-PI31-S168A* ^*attP_ZH-68E*^, *UAS-HA-PI31-S168D* ^*attP_ ZH-68E*^, *HA-PI31*^*Knock-in*^ and *PI31*^*S168A*^ *knock-in* (this study, see below).

Fly genotypes for each figure, table and movie are listed below:

Figure 1A. *w; UAS-HA-PI31/+; R94G06-GAL4/+.*

Figure 1B-G and S1B-E. WT: *yw; +/+; +/+.*

*PI31*^*-/-*^: *w; PI31*^*Δ*^*/PI31*^*Δ*^;*+/+.*

Figure 1H-K and S1F. *yw, hsp70-FLP*^*1*^; *FRT42D, tubP-GAL80/FRT42D, PI31*^*Δ*^; *UAS-IVS-myr::GFP/D42-GAL4.*

Figure 2A and 2B, and Table S2.

Control: *yw; +/+; tubP-GAL4, tubP-GAL80*^*ts*^ */+.*

*HA-PI31*: *w; +/+; tubP-GAL4, tubP-GAL80*^*ts*^*/UAS-HA-PI31-WT* ^*attP_ ZH-68E*^.

Figure 2C and S2E. Control: *yw; +/+; +/+.*

*HA-PI31*^*Knock-in*^: *w; HA-PI31*^*Knock-in*^*/ HA-PI31*^*Knock-in*^; *+/+.*

Figure 3B and Table S3.

*HA-PI31-WT*: *w; +/+; tubP-GAL4, tubP-GAL80*^*ts*^*/UAS-HA-PI31-WT* ^*attP_ ZH-68E*^.

*HA-PI31-S168A*: *w; +/+; tubP-GAL4, tubP-GAL80*^*ts*^*/UAS-HA-PI31-S168A* ^*attP_ ZH-68E*^.

Figure 3C.

*yw*: *w; +/+; tubP-GAL4, tubP-GAL80*^*ts*^*/+.*

*HA-PI31-WT*: *w; +/+; tubP-GAL4, tubP-GAL80*^*ts*^*/UAS-HA-PI31-WT* ^*attP_ ZH-68E*^.

*HA-PI31-S168A*: *w; +/+; tubP-GAL4, tubP-GAL80*^*ts*^*/UAS-HA-PI31-S168A* ^*attP_ ZH-68E*^. *HA-PI31-S168D*: *w; +/+; tubP-GAL4, tubP-GAL80*^*ts*^*/UAS-HA-PI31-S168D* ^*attP_ ZH-68E*^.

Figure 5A and 5B, and Movies S1-3.

WT: *w; +/+; R94G06-GAL4, UAS-Prosβ5-RFP/R94G06-GAL4, UAS-Prosβ5-RFP. PI31*^*-/-*^: *w; PI31*^*Δ*^*/ PI31*^*Δ*^; *R94G06-GAL4, UAS-Prosβ5-RFP/R94G06-GAL4, UAS-Prosβ5-RFP.*

*PI31*^*S168A*^ *knock-in: w; PI31*^*S168A*^*/PI31*^*S168A*^; *R94G06-GAL4, UAS-Prosβ5-RFP/R94G06-GAL4, UAS-Prosβ5-RFP.*

Figure 6A and Table S5. F1 non-Tb flies from the cross of *w; UAS-HA-PI31/UAS-HA-PI31; tubP-GAL4, tubP-GAL80*^*ts*^*/TM6B, Tb* with kinase RNAi transgenic lines.

Figure S1A. *w; +/+; R94G06-GAL4/UAS-mCD8-mCherry.*

Figure S3A. *w; +/+; tubP-GAL4, tubP-GAL80*^*ts*^*/UAS-HA-PI31-WT* ^*attP_ ZH-68E*^.

Figure S4C. *w; PI31*^*S168A*^*/+;+/+.*

Table S1.

*Nrv2-GAL4*: *w;+/+; Nrv2-GAL4, UAS-GFP/Nrv2-GAL4, UAS-GFP.*

*PI31 RNAi*: *w; PI31 RNAi*^*KK105476*^*/PI31 RNAi*^*KK105476*^.

*Nrv2>PI31 RNAi*: *w; PI31 RNAi*^*KK105476*^*/PI31 RNAi*^*KK105476*^; *Nrv2-GAL4, UAS-GFP/Nrv2-GAL4, UAS-GFP.*

### Constructs

The gRNA constructs for making *HA-PI31*^*Knock-in*^ and *PI31*^*S168A*^ *knock-in* fly strains were generated by annealing of two ssDNA oligonucleotides (ssODN) together and inserting the annealed products into the BbsI site of pCFD3-dU6:3gRNA vector (Port et al., 2014) (a gift from Dr. Simon Bullock, Addgene#49410). Listed are the ssODN sequences:

HA-PI31^Knock-in^_F: GTCGAGTCGACCCACTTTCCATTG;

HA-PI31^Knock-in^_R: AAACCAATGGAAAGTGGGTCGACT;

PI31^S168A^ knock-in_F: GTCGTCCAATGGGGCGTGGTGAGT;

PI31^S168A^ knock-in_R: AAACACTCACCACGCCCCATTGGA.

To generate S2 cell vectors to express HA-PI31 and FLAG-dDYNLL1, the HA-PI31 fragment was cut from pcDNA3-HA-PI31(Bader et al., 2011) using EcoRI and XhoI (New England Biolabs), and the FLAG-dDYNLL1 was cloned from a *yw* cDNA library by PCR and the FLAG-tag sequence was added on the forward primer in frame with the dDYNLL1 coding sequence (CDS). HA-PI31 (EcoRI/XhoI) and FLAG-dDYNLL1 (EcoRI/XbaI) fragments were inserted into the S2-cell expression vector pAc5.1/V5-His-A (Thermo Fisher Scientific, V411020).

To generate the S168 point mutation transgenic fly strains (*UAS-HA-PI31-WT* ^*attP_*^ *<SUP>ZH-68E</SUP>, UAS-HA-PI31-S168A* ^*attP_ ZH-68E*^, *UAS-HA-PI31-S168D* ^*attP_ ZH-68E*^), we first mutated the site on pcDNA3-HA-PI31(Bader et al., 2011) through PCR-based site directed mutagenesis. The PCR primers we used are:

S168A_F: CCACGCAGACGACCAACGCCCCACGCCCCATTGGATC;

S168A_R: GATCCAATGGGGCGTGGGGCGTTGGTCGTCTGCGTGG;

S168D_F: CCACGCAGACGACCAACGATCCACGCCCCATTGGATCG;

S168D_R: CGATCCAATGGGGCGTGGATCGTTGGTCGTCTGCGTGG.

After PCR, the template DNA was removed by DpnI treatment. The mutated clones were isolated by Sanger sequencing, and sub-cloned into pUAST-attB vector with EcoRI and XhoI (New England Biolabs)(Bischof et al., 2007).

To generate S2 cell vectors to express HA-PI31 S168 point mutants, the above mutants are sub-cloned into pAc5.1/V5-His-A (Thermo Fisher Scientific, V411020) with EcoRI and XhoI (New England Biolabs).

To generate *E coli.* vectors to recombinantly express different forms of GST-HA-PI31 and FLAG-dDYNLL1, the HA-PI31-WT, -S168A, -S168D and -S168E fragments were cut from pcDNA3-HA-PI31 vectors using EcoRI and XhoI and inserted into the pGEX-4T1 vector (GE Healthcare, #28954549) to generate pGEX-4T1-PI31 clones; the FLAG-dDYNLL1 fragment was produced by PCR using pAc5.1A-FLAG-dDYNLL1 as the template, digested with NcoI and EcoRI (New England Biolabs), and inserted into pET28a vector (Novagen, #69864-3) to generate the pET28a-FLAG-dDYNLL1 clone.

Mouse PI31 (mPI31) CDS was cloned from a cDNA library of mouse small intestine, which was generated by Superscript III First Strand Synthesis Kit (Thermo Fisher Scientific, #18080051) using oligo(dT) amplification, and inserted into the BstI/XmaI sites of pLVX-EF1a-AcGFP-C1 vector (Clontech, #631984). The lentivirus was packaged by transfection of pLVX-EF1a-AcGFP-C1-mPI31, pMD2.G (a gift from Dr. Didier Trono, Addgene #12259) and psPAX2 (a gift from Dr. Didier Trono, Addgene #12260) into HEK293 cells, and collected 48 hours after transfection. Human PI31 (hPI31) CDS was cloned from a cDNA library of HEK293 cells, and inserted into pcDNA3.2/V5/GW/D-TOPO vector (Thermo Fisher Scientific, K244020) through TOPO cloning method (kindly provided by Dr. Park Cho-Park, University of Pennsylvania).

Mouse proteasome subunit α4 CDS was cloned from a mouse cDNA library, and mCherry CDS was subcloned from the mCherry-Mito-7 vector (Olenych et al., 2007) (a gift from Dr. Michael Davidson, Addgene #55102). The α4-mCherry cassette was inserted between the EcoRI/NotI sites of pCAGGS/ES vector. mCerulean CDS was amplified by PCR using mCerulean-N1 vector as the template (Koushik et al., 2006)(a gift from Dr. Steven Vogel, Addgene #27795), and mPI31 CDS was cloned from a mouse cDNA library. The mCerulean-mPI31 cassette was inserted between the EcoRI/NheI sites of pCAGGS/ES vector.

All the constructs were validated by Sanger sequencing (Genewiz).

### Antibody production

To generate a polyclonal antibody against *Drosophila* PI31, GST-PI31 was expressed in BL21 Star (DE3) *E.coli* cells(Thermo Fisher Scientific, #C601003), and purified using Glutathione Sepharose 4B beads (GE Healthcare, #17-0756-01). The purified protein was injected into rabbits and the antisera were collected (Cocalico).

To generate the phosphorylation-specific antibody towards S153 phosphorylation of human PI31, a phospho-peptide was synthesized (Figure S5A), conjugated to carrier proteins and injected into rabbits (YenZym Antibodies, LLC). The elicited antiserum was first affinity-purified against the phospho-peptide. Bound antibodies were eluted and further affinity-absorbed against the non-phosphorylated peptide counterpart to remove antibodies recognizing the other parts of the peptide instead of the phosphorylation site.

### Generation of MARCM clones

*PI31*^*-/-*^ mutant clones were made using the mosaic analysis with a repressible cell marker (MARCM) system(Wu and Luo, 2006). The fly genotype is: *yw, hsp70-FLP*^*1*^; *FRT42D, tubP-GAL80/FRT42D, PI31*^*Δ*^; *UAS-IVS-myr::GFP/D42-GAL4*. The eggs were laid on fly food at 25°C. The 0-6-hour embryos were heat-shocked at 37°C for 30mins followed by recovery at 25 °C for 30mins, and then heat-shocked again at 37°C for 45mins. The wandering 3^rd^-instar larvae with the appropriate genotype were dissected and proceeded for immunostaining.

### Generation of *HA-PI31*^*Knock-in*^ and *PI31*^*S168A*^ *knock-in* fly strains

The off-target effect of gRNAs were assessed by CRISPR target finder (http://tools.flycrispr.molbio.wisc.edu/targetFinder/) (Gratz et al., 2014) and E-CRISPR (www.e-crisp.org/E-CRISP/)(Heigwer et al., 2014). The selected sequences (HA-PI31^Knock-in^:AGTCGACCCACTTTCCATTG, on the “-” strand; PI31^S168A^ knock-in: TCCAATGGGGCGTGGTGAGT, on the “-” strand) were cloned into pCFD3-dU6:3gRNA vector as described above(Port et al., 2014). ssODNs, which serve as donor templates for HDR, were designed with ∼50nt homology arm on each side, and synthesized (IDT). Their sequences are listed below:

HA-PI31Knock-in:

CTTGTACAGCAGATCCCAACCGTAAAAGAAATCGCCCGTCTTGGCAGTCGACCCACTTTCGCCACCGCCGCTAGCATAGTCGGGCACATCATATGGGTACATTGCGGCCACCGATAATCCTTTACAAAATCTGGGTGGGAGTAAACAGAACGAATGG AAC;

PI31^S168A^ knock-in:

GTTCTTACCCGCCACGCCTTGGCTCGCCGATGCGCAAGGGATCTGGATCCGATCCAATGGGGCGTGGTGCGTTCGTCGTCTGCGTGGTAACTTCGCGCGAGTTTCCCGTGAATACAGGGTCTAGAAGCTC.

The pCFD3-dU6:3-PI31 gRNA constructs (final concentration 150ng/µl) and the donor template ssODNs (final concentration 200ng/µl) were mixed together and injected into the *y, sc, v; +/+; nos-Cas9*^*attP2*^ fly (Ren et al., 2013). Knock-in flies were initially identified by PCR and then confirmed by Sanger sequencing (Genewiz).

### Immunofluorescence

Wandering 3^rd^-instar larvae were pinned on a silicone petri dish, and dissected in HL3 saline. The fillet preparations were fixed in 4% paraformaldehyde/PBS for 20mins. After washing with PBS/0.1% Triton X-100 (PBST), the fillets were blocked in 5% NGS/0.2% BSA/PBST, followed by incubation with primary antibodies overnight at 4°C. After washing for 3 times with 0.2%BSA/PBST, the secondary antibodies were applied at room temperature for 3 hours. The fillets were washed 4 times and proceeded for mounting and examination on a confocal microscope (LSM780, Zeiss). The following antibodies were used: chicken anti-mCherry (1:500, Novus, #NBP2-25158), rat anti-HA (1:100, clone 3F10, Sigma-Aldrich, #11867423001), mouse anti-ubiquitin conjugates (1:500, clone FK2, Enzo, #PW8810), mouse anti-Brp (1:300, clone nc82, Developmental Studies Hybridoma Bank, #nc82), chicken anti-GFP (1:1000, Aves Labs, #GFP-1020), rabbit anti-GluRIIC(Marrus et al., 2004) (1:1000, kindly provided by Dr. Aaron DiAntonio, Washington University in St. Louis), goat anti-rat-Alexa 488(1:500, Jackson ImmunoResearch, #112-545-167), goat anti-mouse-Alexa 488 plus (1:500, Thermo Fisher Scientific, #A32723), goat anti-chicken-Alexa 488(1:500, Jackson ImmunoResearch, #103-545-155), goat anti-chicken-Cy3(1:500, Jackson ImmunoResearch, #103-165-155), goat anti-mouse-Alexa 568(1:500, Thermo Fisher Scientific, #A11031), goat anti-HRP-Cy3(1:3000, Jackson ImmunoResearch, #123-165-021), goat anti-rabbit-Alexa 633(1:500, Thermo Fisher Scientific, #A21071), and goat anti-HRP-Alexa 647(1:300, Jackson ImmunoResearch, #123-605-021). Quantification of puncta number and intensity of Poly-Ub proteins was done with Fiji software(Schindelin et al., 2012), and Fiji and Imaris (Bitplane) softwares were used for quantification of numbers and volume of Brp puncta.

### Co-Immunoprecipitation (Co-IP)

For the Co-IP experiments shown in Figure 2A, 2B, 3B, and 3C, *tubP-GAL4, tubP-GAL80*^*ts*^ flies were crossed with indicated forms of *UAS-HA-PI31* or *yw* (as control) flies and raised at 25°C. The 0-3-day-old F1 progenies were collected and moved to 29°C for 6 days. Three hundred male flies were collected and homogenized in 1.5ml buffer A [50mM Tris-HCl (pH7.5), 137mM NaCl, 0.25% NP-40, protease inhibitor cocktail (Roche, #11873580001) and phosphatase inhibitor cocktail (Roche, #04906837001)]. After centrifuged at 20000g for 15mins and filtered through a low-protein binding 0.45µm PVDF membrane syringe filter (Millipore, #SLHVX13NL), the extracts were incubated with 30µl mouse anti-HA agarose beads (clone HA-7, Sigma-Aldrich, #A2095) at 4°C for 3 hours. After six washes in buffer A and two additional washes in buffer A without NP-40 and protease and phosphatase inhibitors, the beads were eluted in 30µl 0.1M glycine (pH2.5) at room temperature for 10mins. The eluted proteins were taken either for mass spectrometry (MS) analysis (Figure 2A and 3B) or for Western Blot analysis (Figure 2B and 3C).

For the Co-IP experiment shown in Figure 2C, 150 3-day old male *HA-PI31*^*Knock-in*^ flies and *yw* flies (as control) were collected separately and homogenized in 1ml buffer A with 50U/ml DNase I (Thermo Fisher Scientific, #18047019). After centrifuged at 20000g for 15mins, sonicated briefly to break genomic DNAs, and filtered through a 0.45µm syringe filter, the extracts were incubated with 30µl mouse anti-HA magnetic beads (Thermo Fisher Scientific, #88836) at 4°C for 3 hours. After five washes in buffer A and two additional washes in buffer A without NP-40 and protease and phosphatase inhibitors, the beads were eluted in 30µl 500µg/ml HA peptide (Thermo Fisher Scientific, #26184) at 37°C for 30mins. The eluted proteins were analyzed by Western blotting.

For the IP or Co-IP experiments shown in Figure 2D, 2E, 4B, 4C, S2C, S2D, S3C and S3D, 10cm-plate cultures of indicated cell lines were transfected/infected with indicated constructs using TransIT-Insect transfection reagent (Mirus, #MIR6100) for S2 cells, Lipofectamine 2000 transfection reagent (Thermo Fisher Scientific, #11668-019) for HEK293 cells and lentivirus for primary MEFs, respectively. After 48 hours for the transfected cells or 7 days for the infected MEFs, the cell cultures were rinsed in cold PBS, collected, and resuspended in 0.6ml buffer A. Lysis was facilitated by freeze/thaw for three times. After centrifugation at 20000g for 15mins and filtration through a 0.45µm syringe filter, the extracts were incubated with 30µl mouse anti-HA agarose beads (clone HA-7, Sigma-Aldrich, #A2095), or 30µl mouse anti-FLAG magnetic beads (clone M2, Sigma-Aldrich, #M8823), or 30µl GFP-Trap agarose beads (Chromotek, gta-20) at 4°C for 3 hours. The beads were washed in buffer A for six times and then were eluted in 30µl 1.5XSDS-PAGE sample buffer by boiling at 95°C for 5mins.

For the Co-IP experiment shown in Figure 4A, 10-cm plate cultures of S2 cells were transected with 15µg pAc5.1A-FLAG-dDYNLL1 or pAc5.1A empty vector (as control) constructs using TransIT-Insect transfection reagent. After 48 hours, the cells were rinsed, collected and resuspended in 600µl buffer A, and lysis was facilitated by freeze/thaw for three times. After centrifugation at 20000g for 15mins and filtration through a 0.45µm syringe filter, the extracts were incubated with 30µl mouse anti-FLAG magnetic beads (clone M2, Sigma-Aldrich, #M8823) at 4°C for 3 hours. After six washes in buffer A and two additional washes in buffer A without NP-40 and protease and phosphatase inhibitors, the beads were eluted in 30µl 300 µg/ml 3XFLAG peptide (Sigma-Aldrich, #F4799) at 4°C for 45mins. The eluted proteins were run on a SDS-PAGE gel for ∼1cm to remove the 3XFLAG peptide. Finally, the gel was cut for MS analysis. Biological triplicates of the Co-IP were used for quantification.

For the Co-IP experiment shown in Figure S2E, eight hundreds of 3-day-old male *HA-PI31*^*Knock-in*^ flies and *yw* flies (as control) were collected separately and homogenized in 5ml buffer B [25mM HEPES (pH7.5), 100mM NaCl, 5mM MgCl2, 2mM ATP, 10% glycerol, 0.25% NP-40, protease inhibitor cocktail and phosphatase inhibitor cocktail] with 50U/ml DNase I (Thermo Fisher Scientific, #18047019). After centrifugation at 20000g for 15mins, lysates were sonicated briefly to break genomic DNA and filtered through a 0.45µm syringe filter, and then the extracts were incubated with 200µl mouse anti-HA magnetic beads (Thermo Fisher Scientific, #88836) at 4°C for 3 hours. After five washes in buffer B and two additional washes in buffer C [25mM HEPES (pH7.5), 150mM NaCl, 5mM MgCl_2_, 2mM ATP, 10% glycerol], the beads were eluted by 200µl 500µg/ml HA peptide in buffer C at 37°C for 30mins. The eluted proteins were concentrated 5X with 10KDa-cutoff Amicon centrifugal filter unit (Millipore, #UFC501024), and analyzed by native gels and Western blotting.

### Mass spectrometry analysis

Glycine (0.1M, pH2.5) eluted samples (Figure 2A, 3B, 6A, S3A and S3B) were dried in Speedvac concentrator (Thermo Fisher Scientific) and dissolved in 8 M urea, 0.1 M ammonium bicarbonate, and 10 mM DTT. After reduction, cysteines were alkylated in 30mM iodoacetamide. The proteins were digested in less than 4 M urea by endoproteinase LysC (Wako Chemicals) followed by digestion with Trypsin Gold (Promega) in less than 2 M urea. FLAG-peptide eluted samples (Figure 4A) were cleaned up using SDS-PAGE, reduced (10 mM DTT) and alkylated (30 mM iodoacetamide), followed by digestion with a cocktail of LysC and Trypsin Gold. Digestions were halted by adding trifluoroacetic acid (TFA) and digests were desalted and analyzed by reversed phase nano-LC-MS/MS using either a Fusion Lumos or a Q-Exactive Plus (Thermo Fisher Scientific) both operated in high/high mode(Rappsilber et al., 2007).

Data were searched and quantified against Uniprot’s *Drosophila* and human proteome databases using ProteomeDiscoverer v. 1.4.0.288 (Thermo Scientific) combined with Mascot v. 2.5.1 (Matrix Science) and/or MaxQuant v. 1.6.0.13. (Cox et al., 2014). Oxidation of methionine and protein N-terminal acetylation were allowed as variable modifications and all cysteines were treated as being carbamidomethylated. Peptide matches were filtered using a Percolator-calculated false discovery rate (FDR) of 1%(Kall et al., 2007). For the quantitative MaxQuant analysis, FDR thresholds for peptides and proteins were set at 2% and 1%, respectively. For assignment of phosphorylated residues, we used PhosRS3.0 combined with Mascot(Taus et al., 2011).

Label Free Quantitation (LFQ) or intensity-based absolute quantification (iBAQ) were used for the identification and quantitation of binding partners(Cox et al., 2014; Schwanhausser et al., 2011). Data were processed using Perseus v1.6.0.7(Tyanova et al., 2016). In short, reverse matches and common contaminations were disregarded, and protein signals in minimum 2 out of 3 replicates for at least one condition was required. Data were validated by scatter plots, LFQ or iBAQ based principal component analysis (PCA) and distributions of LFQ or iBAQ values, and found to be comparable. T-tests were performed and proteins were considered as interesting hits if showing a difference of more than 4-linear folds and a p-value of less than 0.01.

### GST pull-down assays

BL21 Star (DE3) competent *E.coli* cells were transformed with pGEX-4T1-PI31 vectors (-WT, -S168A, -S168D, and -S168E), pGEX-4T1 empty vector or pET28a-FLAG-dDYNLL1 vector. Single colonies were picked and cultured in 5ml LB, followed by an enlarged culture in 100ml LB. When OD_600_≈0.4, IPTG was added to a final concentration of 0.3mM. After 4 hours, the bacteria were collected by centrifugation and rinsed by cold PBS. The bacteria were resuspended in 5ml cold PBS with protease inhibitor cocktail, lysed by sonication and centrifuged at 20000g for 15mins.

1ml pGEX-4T1 empty vector lysate or 1ml pGEX-4T1-PI31 lysate were incubated with 30µl Glutathione Sepharose 4B beads at 4°C for 3 hours. Then, the beads were washed with PBS for three times, followed by incubation with 1ml pET28a-FLAG-dDYNLL1 lysate at 4°C for 3 hours. The beads were washed in buffer A for five times and in 50 mM Tris-HCl (pH 8.0) once. Elution was done by incubation of the beads in 30µl GST elution buffer [50 mM Tris-HCl (pH 8.0) and 20mM reduced glutathione (Sigma-Aldrich, #G4251)] at room temperature for 20mins. The eluted proteins were analyzed by SDS-PAGE and Western blot. Ponceau S (Sigma-Aldrich, #P7170) staining was used to assess purity and amount of GST and different forms of GST-PI31 proteins. Anti-FLAG-HRP (1:1000, Sigma-Aldrich, #A8592) was used to evaluate amount of FLAG-dDYNLL1 bound to GST or different forms of GST-PI31 proteins.

### *In vitro* reconstitution of dDYNLL1-PI31-proteasome complex

Six-milliliter lysate of BL21 Star (DE3) expressing pET28a-FLAG-dDYNLL1 was incubated with 200µl mouse anti-FLAG magnetic beads (clone M2, Sigma-Aldrich, #M8823) at 4°C for 3 hours. After washed with buffer A for four times, the beads were split equally into six tubes. Then, 5µg purified bovine 20S proteasomes (UBPbio, #A1401), or 5µg GST+5µg purified bovine 20S proteasomes, or 5µg indicated forms of GST-PI31 proteins+5µg purified bovine 20S proteasomes were added to the six tubes, respectively, and incubated at 4°C for 3 hours. After four washes in buffer A, the beads were eluted in 35µl 300µg/ml 3XFLAG peptide at 4°C for 45mins.

### Kinase screen

The conserved PI31 phosphorylation site (pSerine-Proline) pointed to Proline-directed kinases, such as cyclin-dependent kinases (CDKs) and mitogen-activated protein kinases (MAPKs) (Wagih et al., 2016; Zhu et al., 2005). Therefore, we screened all available CDK and MAPK RNAi fly strains from the Transgenic RNAi Project (TRiP) collection. *Tub-GAL4, Tub-GAL80*^*ts*^*>UAS-HA-PI31* flies were crossed with RNAi transgenic fly strains. HA-PI31 protein was immunoprecipitated and digested. The peptides containing the phosphorylated and non-phosphorylated S168 (EVTTQTTN^168^SPRPIGSDPDPLR) were targeted and measured in parallel reaction monitoring (PRM) experiments. Phospho-peptide signal versus total signal was used to estimate the phosphorylation occupancy percentage.

### *In vitro* kinase assay

Purified recombinant human p38α (MAPK14, GST-tagged, Thermo Fisher Scientific, #PV3304), p38β (MAPK11, His-tagged, Thermo Fisher Scientific, #PV3679), p38γ (MAPK12, His-tagged Thermo Fisher Scientific, #PV3654), p38δ (MAPK13, His-tagged, Thermo Fisher Scientific, #PV3656) and hPI31 (UBPBio, #A3901) were used for *in vitro* kinase assay.

One microgram of hPI31 was incubated with 200ng indicated kinases in 30µl kinase assay buffer [20 mM Tris (pH 7.5), 2 mM DTT, 10 mM MgCl2, 200µM ATP, and phosphatase inhibitors] at 30°C for 30mins. The reaction was terminated by adding 15µl 3XSDS-PAGE loading buffer and boiling at 95°C for 5mins.

### *In vitro* phosphatase assay

To validate the specificity of p153-hPI31 antibody, HA-hPI31 was expressed in HEK293 cells and IP with anti-HA agarose beads. After four washes, the beads were split into three tubes. One was incubated with 50µl phosphatase assay buffer (50mM HEPES, 100mM NaCl, 2mM DTT, 0.01% Brij-35) as a control, one was treated with 400U Lambda protein phosphatase (Lambda PP, New England Biolabs, #P0753S) in 50µl phosphatase assay buffer, and the other one was treated with Lambda PP in the presence of phosphatase inhibitor cocktail (Roche, #04906837001). After incubated at 30°C for 30mins, the beads were precipitated and hPI31 proteins were eluted in 1XSDS-PAGE loading buffer and boiled at 95°C for 5mins.

### Native gel assays

Protein complexes captured by anti-HA Co-IP using extracts of *yw* (as control) and *HA-PI31*^*Knock-in*^ flies were eluted in buffer C with HA peptide under the native condition (see the “Co-IP” section for more details). The eluted proteins, as well as 15µg total lysates of *yw* and *HA-PI31*^*Knock-in*^ flies (as inputs for the Co-IP experiment), were mixed with 2X native sample buffer (Bio-rad, #1610738), and loaded onto 3-8% Tris-acetate gels (Bio-rad, #3450131). Purified bovine 19S (UBPbio, #A1301), 20S (UBPbio, #A1401) and 26S (UBPbio, #A1201) proteasomes were also loaded onto the gels as standards to mark positions of different proteasome complexes. Electrophoresis was started with 50V for 1 hour in running buffer (90mM Tris, 90mM Boric Acid, 1mM EDTA, 2.5 mM MgCl2, 1mM ATP and 0.5mM DTT). After samples fully entered the gel, voltage was increased to 120V and ran 5 hours at 4°C. Then, proteins in the gels were transferred to PVDF membrane for Western Blot.

### Western Blot Analysis

Proteins in SDS-PAGE gels or native gels were transferred onto 0.45µm Immobilon-P PVDF membranes (Millipore. #IPVH00010). The membranes were blocked with 5% non-fat milk in TBST [20mM Tris (pH7.5), 300mM NaCl, 0.1% Tween-20] at room temperature for 1 hour, followed by incubation with primary antibodies in 5% BSA/TBST at 4°C overnight. After washed for three times with TBST, the membranes were incubated with HRP-conjugated secondary antibodies at room temperature for 1.5 hours. The membranes were washed for four times with TBST, and detection was performed with ECL reagent (GE Healthcare, #RPN2134) or SuperSignal West Femto Maximum Sensitivity Substrate (Thermo Fisher Scientific, # 34096). The following antibodies were used for Western blot: rabbit anti-DYNLL1/2(1:3000, clone EP1660Y, Abcam, #ab51603), mouse anti-α7 (1:3000, clone MCP72, Enzo, # PW8110), mouse anti-Rpt3 (1:3000, clone TBP7-27, Enzo, # PW8765), rabbit anti-dPI31(1:5000), goat anti-hPI31(1:3000, Thermo Fisher Scientific, #PA5-18133), rabbit anti-hPI31(1:3000, Abcam, #ab140497), rabbit anti-phospho-p38 MAPK (Thr180/Tyr182) (1:3000, clone 3D7, Cell Signaling Technology, #9215), rabbit anti-p38 MAPK (total) (1:3000, Cell Signaling Technology, #9212), rabbit anti-6X His-tag (1:3000, Abcam, #ab9108), rat anti-HA-HRP(1:3000, clone 3F10, Roche, #12013819001), mouse anti-FLAG-HRP(1:1000, clone M2, Sigma-Aldrich, #A8592), mouse anti-GFP-HRP(1:1000, clone B2, Santa Cruz Biotechnology, #sc-9996-HRP), rabbit anti-GST-HRP(1:1000, Abcam, #ab3416), rabbit anti-β-Actin-HRP (1:10000, clone 13E5, Cell Signaling Technology, #5125), donkey anti-rabbit-HRP(1:5000, Jackson ImmunoResearch, #711-035-152), donkey anti-mouse-HRP(1:5000, Jackson ImmunoResearch, #715-035-150), goat anti-mouse Fc-HRP(1:3000, Thermo Fisher Scientific, #31437).

### Live imaging

For analysis of proteasome transport in axons of *Drosophila* motor neurons, wandering 3^rd^-instar larvae of *R94G06-GAL4>Prosβ5-RFP* (as control), *PI31*^*-/-*^, *R94G06-GAL4>Prosβ5-RFP and PI31*^*S168A*^, *R94G06-GAL4>Prosβ5-RFP* were dissected in HL3 saline supplemented with 7mM L-glutamic acid, and flipped the inside out to expose motor neuron axons. The larval preparations were mounted on a microscope slide with coverslip bridges to avoid crash of tissues. Live imaging was done on a 60X silicone oil objective lens (NA1.3, Olympus) of inverted Olympus IX-70 microscope equipped with Evolve-512 EMCCD camera (Photometrics). Images were captured using a 500-ms exposure at 1.4 Hz, and each recording consisted of 100 frames in 70 seconds. Kymographs were generated using ImageJ with the “Multiple Kymograph” plugin. All particles were picked and tracked by the software.

For live imaging of α4-mCherry and mCerulean–PI31 in mouse DRG neuron cultures, experiments were approved by Animal Care and Use Committee (IACUC) of The Rockefeller University (protocol number 17006). DRGs were dissected from E13 mouse embryos (CD-1 IGS pregnant mouse, Charles River, #022) and dissociated to single cell suspension using Trypsin. Dissociated sensory neurons were transfected with pCAGGS/ES-α4-mCherry and pCAGGS/ES-mCerulean–mPI31 plasmids using the Neon transfection system (Thermo Fisher Scientific, MPK5000) at 1400 V, 20 ms, and 1 pulse. After transfection, 5X10^5^ neurons were centrifuged, resuspended in 9µl growth medium [Neurobasal medium (Thermo Fisher Scientific, #21103049), B-27 (Thermo Fisher Scientific, #17504044), penicillin-streptomycin (Thermo Fisher Scientific, #15140122) with 12.5ng/ml mouse growth factor 2.5S (NGF, Alomone Labs, #N-100)] and transferred to 15µl growth medium as a hanging drop overnight to form cell aggregates. Aggregates were cut into four pieces, plated on Poly-D-Lysine (Sigma-Aldrich, #P6407)/Laminin (Sigma-Aldrich, #L2020)-coated 8-well µ-Slide (ibidi, # 80826) and cultured for additional 24-48 hours. Before imaging, culture medium was changed to Hibernate E low fluorescence media (BrainBits, #HELF) supplemented with B27 and NGF, in order to maintain cells in ambient CO_2_ levels. Images were taken on a 60X silicone oil objective lens (NA1.3, Olympus) of inverted Olympus IX-70 microscope equipped with Evolve-512 EMCCD camera (Photometrics) and with a heating chamber at 37°C, at 0.5Hz for 1-3 minutes.

## Supplementary Information

### Supplementary Information includes five figures, four movies and five tables. Supplementary Figure Legends

**Figure S1.**
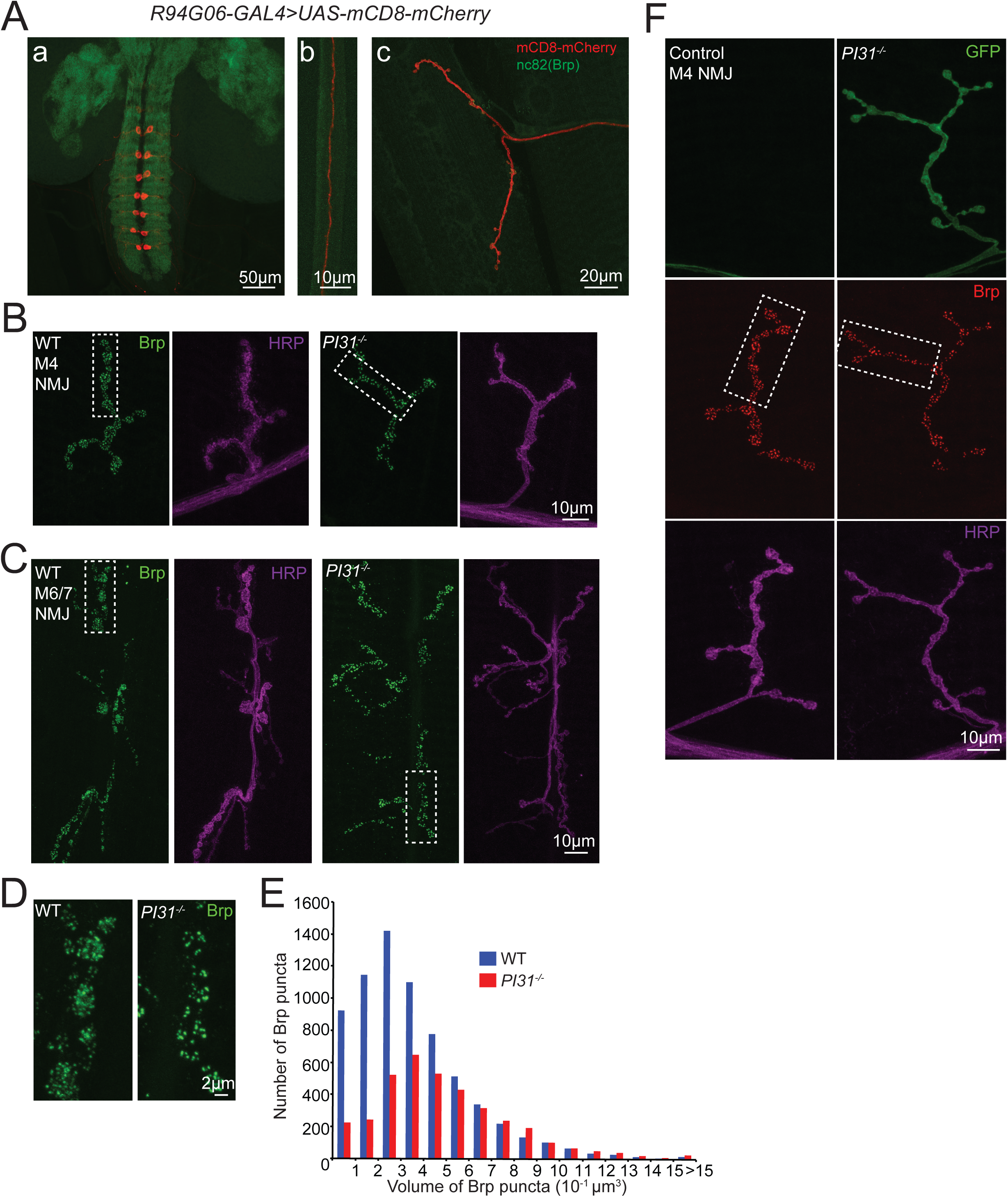
Structural changes of synapses at neuromuscular junctions (NMJs) of *PI31* ^*-/-*^ larva. Related to Figure 1. **(A) Expression pattern of *R94G06-GAL4* illustrated by using it to drive *UAS-mCD8-mCherry*. (a)** Brain and ventral nerve cord; **(b)** A single axon in nerve; **(c)** A NMJ at Muscle 1. Brp antibody (clone nc82, green) was used to visualize the synaptic neuropil regions. The results indicate that *R94G06-GAL4* is a specific driver for a subset of motor neurons (MN1-IB) targeting Muscle 1. **(B)** Overview images for M4 NMJs of wild-type and *PI31*^*-/-*^ larvae immunostained with anti-Brp (green) and anti-HRP (magenta) antibodies. They indicate that the overall morphology of NMJs is not dramatically changed, but the number of Brp puncta is largely reduced in *PI31* **^*-/-*^** larvae, suggesting a structural defect of synapses. High-magnification images of the boxed area are shown in Figure 1F, and quantification of Brp puncta is shown in Figure 1G. **(C and D)** Representative images of M6/7 NMJs indicate that the number of Brp puncta is largely reduced in *PI31* **^*-/-*^** larvae. **(C)** Overview images of M6/7 NMJs. **(D)** High-magnification images of the boxed area in (C). See quantification of Brp puncta in Figure 1G. **(E)** Frequency distribution plot of Brp punctum volumes shows that small Brp puncta (volume<0.5µm^3^) were specifically affected by loss of PI31. The images used for the quantification of Brp punctum number shown in Figure 1G were analyzed here for Brp punctum volumes. n=6853 Brp puncta from 20 M4 NMJs for wild-type, and n= 3672 Brp puncta from 20 M4 NMJs for *PI31*^*-/-*^. **(F)** Analysis of *PI31*^*-/-*^ MARCM clones reveals that the reduction of Brp puncta is a cell-autonomous phenotype. The GFP-positive neuron is *PI31*^*-/-*^, whereas the GFP-negative neuron is *PI31*^*+/-*^ (control) from the same larva. High-magnification images of the boxed regions are shown in Figure 1H, and quantification of Brp puncta is shown in Figure 1I.

**Figure S2.**
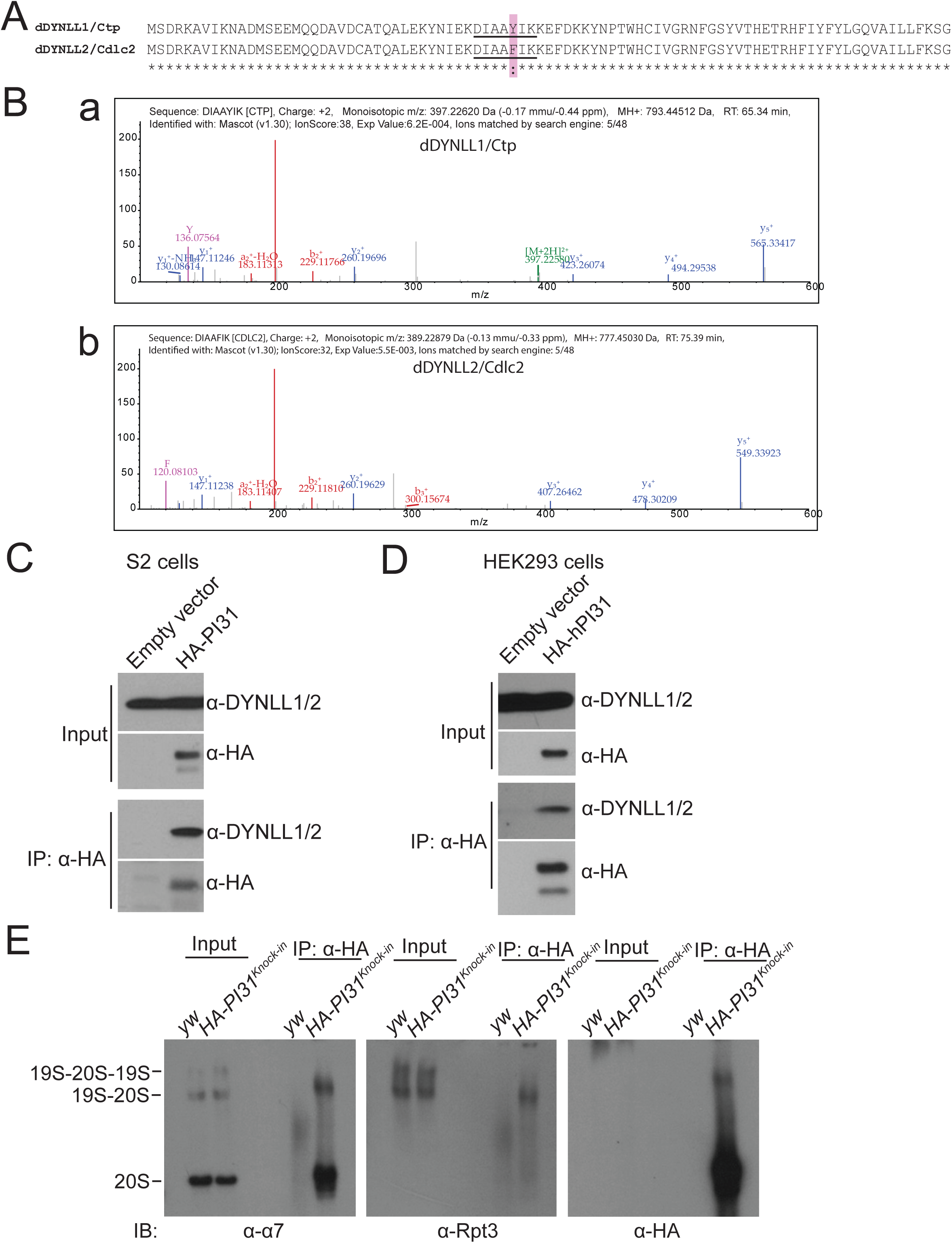
DYNLL1/2 are binding partners of PI31. Related to Figure 2. **(A)** Alignment of dDYNLL1/Ctp and dDYNLL2/Cdlc2. They differ in only one amino acid, which is highlighted by a pink box. The alignment was performed with Clustal Omega program on Uniprot server. **(B)** The spectra of the unique peptides for dDYNLL1/Ctp (**a**, sequence: DIAAYIK) and dDYNLL2/Cdlc2 (**b**, sequence: DIAAFIK). The peptides were identified in the anti-HA-PI31 Co-IP samples using nano liquid chromatography and high-resolution MS. Both of them were matched with high confidence and with mass accuracy better than 0.5ppm. These results indicate that both proteins interact with PI31. **(C)** Western blot analysis of anti-HA-PI31 Co-IP experiments in S2 cells indicates that PI31 interacts dDYNLL1/2. **(D)** Western blot analysis of anti-HA-hPI31 Co-IP experiments in HEK293 cells indicates that the interaction between PI31 and DYNLL1/2 is conserved in human cells. **(E)** Native gel-Western blot analysis of PI31-bound complexes reveals that PI31 binds to both 20S and single-capped 26S proteasomes. *HA-PI31*^*Knock-in*^ flies were collected for anti-HA Co-IP experiments. HA-PI31 and its interacting proteins were eluted by HA peptide under native conditions, and resolved on native gels. ATP was present during the Co-IP and native gel procedures. Anti-α7 and anti-Rpt3 antibodies were used for Western blot analysis to visualize 20S(α7), 19S(Rpt3) and 26S(both α7 and Rpt3) proteasomes.

**Figure S3.**
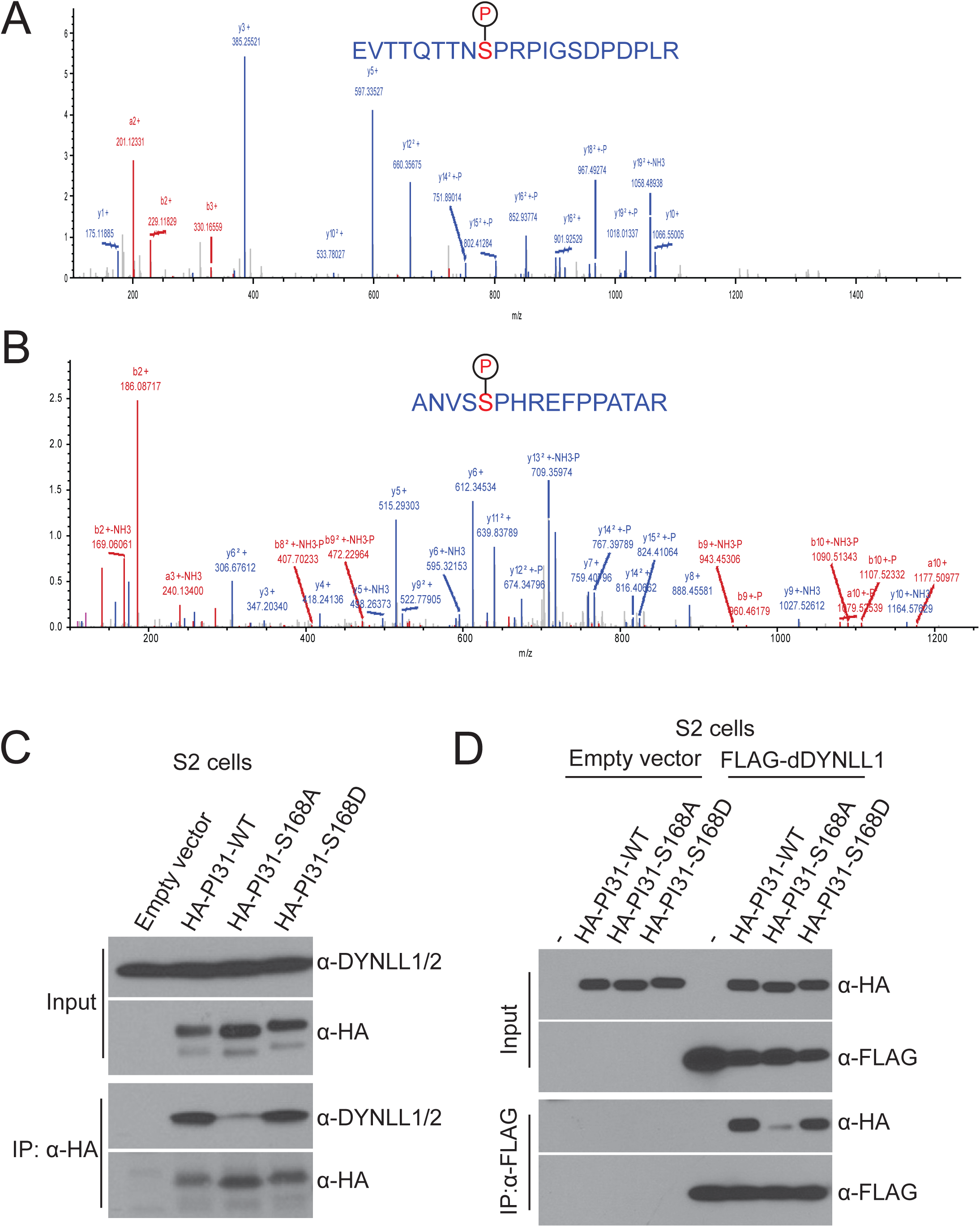
PI31 phosphorylation regulates interaction with dDYNLL1. Related to Figure 3. **(A and B)** High resolution/high mass accuracy tandem MS spectra for the phospho-peptides of *Drosophila* **(A)** and human **(B)** PI31. Both peptides were matched with mass accuracy of <1ppm. Peptide information is shown below and probabilities for phosphorylated residues, calculated using PhosphoRS 3.0, are show in square brackets. *Drosophila*, EVTTQTTNSPRPIGSDPDPLR [T(3): 0.0; T(4): 0.0; T(6): 0.0; T(7): 0.0; S(9): 100.0; S(15): 0.0], Charge: +2, Monoisotopic m/z: 1181.05811; Human, ANVSSPHREFPPATAR [S(4): 0.5; S(5): 99.5; T(14): 0.0], Charge: +3, Monoisotopic m/z: 606.28796. **(C)**Co-IP-Western blot analysis shows that the non-phosphorable mutation of PI31 (S168A) largely diminished interaction with dDYNLL1/2 in S2 cells. **(D)**Reciprocal Co-IP using FLAG-dDYNLL1 as the bait confirmed that the PI31-dDYNLL1 interaction was phosphorylation-dependent.

**Figure S4.**
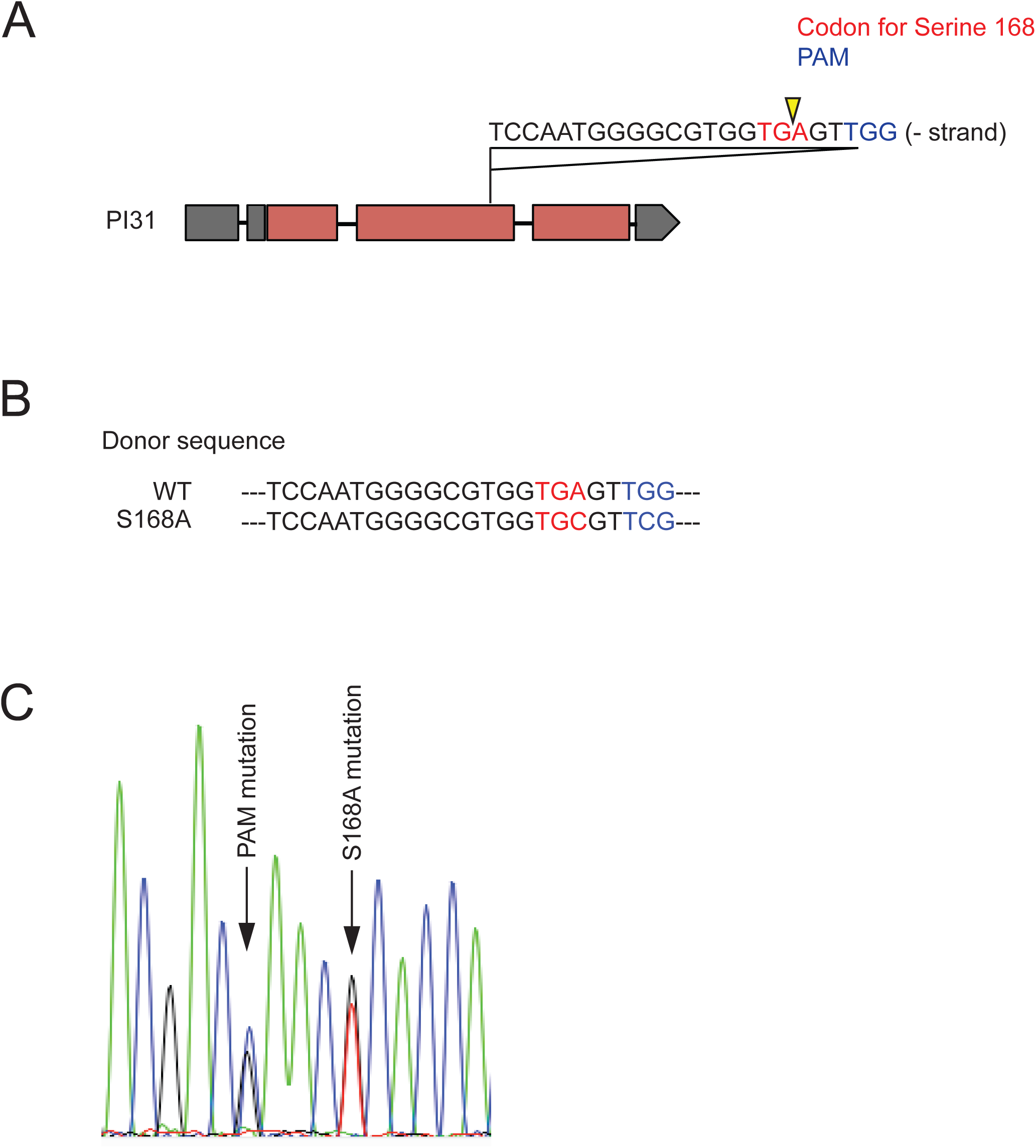
Generation of *PI31*^*S168A*^ *knock-in* fly strain. Related to Figure 5. **(A)** Schematic diagram shows the sequence and position of the gRNA target site in the PI31 locus. UTRs are shown as gray blocks, and coding sequences are shown as dark red blocks. The codon for S168 is shown in red, whereas the PAM sequence is shown in blue. The Cas9 cut site is indicated by a yellow arrowhead. **(B)** Partial sequence of the HDR donor DNA used to generate the S168A mutant. The codon for S168 is shown in red, whereas the PAM sequence is shown in blue. In order to avoid repeated cut by Cas9, we also introduced a silent mutation to disrupt the PAM. **(C)** The chromatogram obtained through Sanger sequencing of heterozygous *S168A knock-in* flies. The arrows point out positions of the S168A and PAM mutations. Double peaks represent an overlay of the sequences from the S168A mutant and wild-type PI31 alleles. This result verified the precise integration of the donor DNA to the PI31 locus.

**Figure S5.**
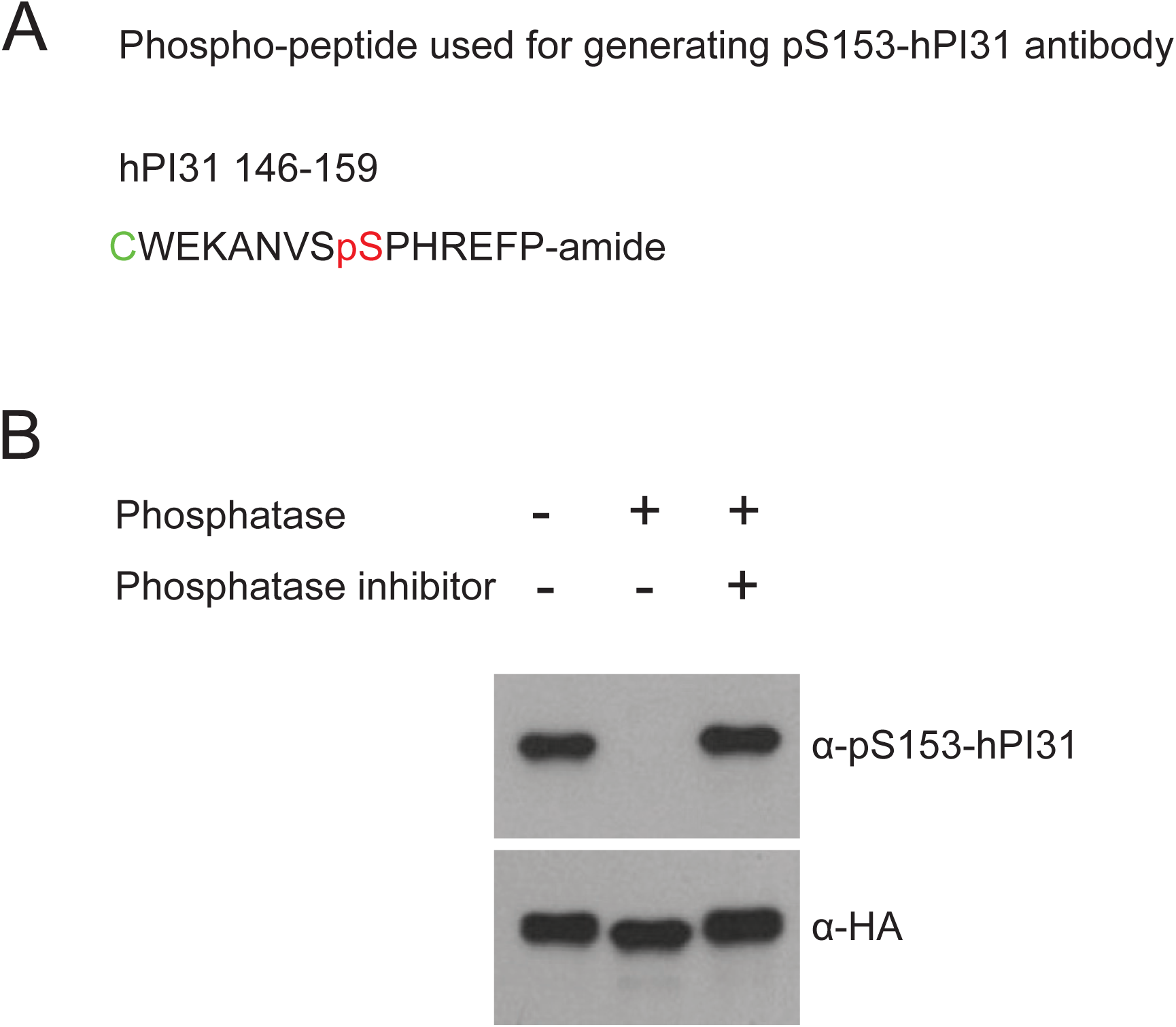
Generation and validation of pS153-hPI31 antibody. Related to Figure 6. **(A)** Sequence of the phospho-peptide used for the generation of pS153-hPI31 antibody. The phosphorylated residue is in red, and the N-terminal cysteine that is in green was added to facilitate conjugation of the peptide to the carrier protein for immunization. **(B)** Validation of the antibody specificity by *in vitro* phosphatase assay. HA-hPI31 expressed in HEK293 cells was immunoprecipitated by anti-HA agarose beads and treated with 400U lambda phosphatase at 30°C for 30 minutes with or without phosphatase inhibitors. The results show that the antibody recognized a clear band around 31kD, which was abolished by the phosphatase but rescued back by the phosphatase inhibitors. This data demonstrates that the pS153-hPI31 antibody specifically recognizes the phosphorylated hPI31.

### Supplementary Movie Legends

**Movie S1. A representative video of Prosβ5-RFP particle movement in an axon of wild-type larva (*w; +/+; R94G06-GAL4, UAS-Prosβ5-RFP/R94G06-GAL4, UAS-Prosβ5-RFP*). Related to Figure 5.** The video was acquired at 1.4Hz rate for 70 seconds (100 frames in total), and is shown in 10X of the original speed.

**Movie S2. A representative video of Prosβ5-RFP particle movement in an axon of *PI31*^*-/-*^ larva (*w; PI31*^*Δ*^*/ PI31*^*Δ*^; *R94G06-GAL4, UAS-Prosβ5-RFP/R94G06-GAL4, UAS-Prosβ5-RFP*). Related to Figure 5.** The video was acquired at 1.4Hz rate for 70 seconds (100 frames in total), and is shown in 10X of the original speed.

**Movie S3. A representative video of Prosβ5-RFP particle movement in an axon of *PI31*^*S168A*^ *knock-in* larva (*w; PI31*^*S168A*^ */ PI31*^*S168A*^; *R94G06-GAL4, UAS-Prosβ5-RFP/R94G06-GAL4, UAS-Prosβ5-RFP*). Related to Figure 5.** The video was acquired at 1.4Hz rate for 70 seconds (100 frames in total), and is shown in 10X of the original speed.

**Movie S4**. **A representative video of mCerulean-mPI31 (green) and α4-mCherry (red) movement in axons of cultured mouse DRG neurons. Related to Figure 5.** The video was acquired at 0.5Hz rate for 140 seconds (70 frames in total), and is shown in 10X of the original speed.

### Supplementary Table Legends

**Table S1.**
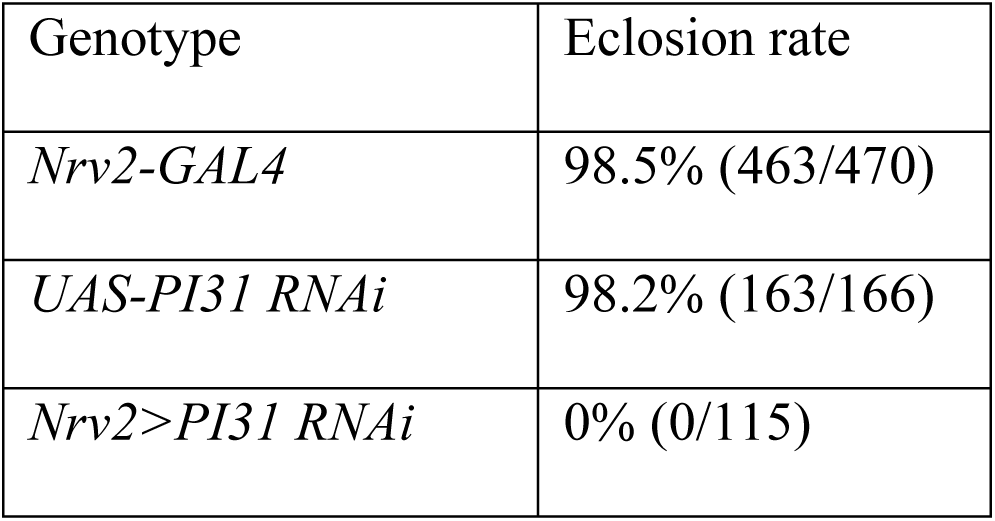
**Inactivation of PI31 in neurons results in an eclosion defect. Related to Figure 1.**

**Table S2. Raw data for the MS analysis of the anti-HA-PI31 Co-IP experiments. Related to Figure 2.**

**Table S3. Raw data for the MS analysis of the anti-HA-PI31^S168A^ Co-IP versus anti-HA-PI31^WT^ Co-IP experiments. Related to Figure 3.**

**Table S4. Raw data for the MS analysis of the anti-FLAG-dDYNLL1 Co-IP experiments. Related to Figure 4.**

**Table S5. Summary for the MS analysis of the kinase screen experiments. Related to Figure 6.**

## References

Asano, S., Fukuda, Y., Beck, F., Aufderheide, A., Forster, F., Danev, R., and Baumeister, W. (2015). Proteasomes. A molecular census of 26S proteasomes in intact neurons. Science 347, 439–442.

Bader, M., Benjamin, S., Wapinski, O.L., Smith, D.M., Goldberg, A.L., and Steller, H. (2011). A conserved F box regulatory complex controls proteasome activity in Drosophila. Cell 145, 371–382.

Bai, C., Sen, P., Hofmann, K., Ma, L., Goebl, M., Harper, J.W., and Elledge, S.J. (1996). SKP1 connects cell cycle regulators to the ubiquitin proteolysis machinery through a novel motif, the F-box. Cell 86, 263–274.

Balch, W.E., Morimoto, R.I., Dillin, A., and Kelly, J.W. (2008). Adapting proteostasis for disease intervention. Science 319, 916–919.

Ballatore, C., Lee, V.M., and Trojanowski, J.Q. (2007). Tau-mediated neurodegeneration in Alzheimer’s disease and related disorders. Nat Rev Neurosci 8, 663–672.

Baumeister, W., Walz, J., Zuhl, F., and Seemuller, E. (1998). The proteasome: paradigm of a self-compartmentalizing protease. Cell 92, 367–380.

Bento, C.F., Renna, M., Ghislat, G., Puri, C., Ashkenazi, A., Vicinanza, M., Menzies, F.M., and Rubinsztein, D.C. (2016). Mammalian Autophagy: How Does It Work? Annual review of biochemistry 85, 685–713.

Bilen, J., and Bonini, N.M. (2005). Drosophila as a model for human neurodegenerative disease. Annual review of genetics 39, 153–171.

Bingol, B., and Schuman, E.M. (2006). Activity-dependent dynamics and sequestration of proteasomes in dendritic spines. Nature 441, 1144–1148.

Bingol, B., and Sheng, M. (2011). Deconstruction for reconstruction: the role of proteolysis in neural plasticity and disease. Neuron 69, 22–32.

Bischof, J., Maeda, R.K., Hediger, M., Karch, F., and Basler, K. (2007). An optimized transgenesis system for Drosophila using germ-line-specific phiC31 integrases. Proceedings of the National Academy of Sciences of the United States of America 104, 3312–3317.

Campbell, D.S., and Holt, C.E. (2001). Chemotropic responses of retinal growth cones mediated by rapid local protein synthesis and degradation. Neuron 32, 1013–1026.

Chintapalli, V.R., Wang, J., and Dow, J.A. (2007). Using FlyAtlas to identify better Drosophila melanogaster models of human disease. Nature genetics 39, 715–720.

Cho-Park, P.F., and Steller, H. (2013). Proteasome regulation by ADP-ribosylation. Cell 153, 614–627.

Chu-Ping, M., Slaughter, C.A., and DeMartino, G.N. (1992). Purification and characterization of a protein inhibitor of the 20S proteasome (macropain). Biochimica et biophysica acta 1119, 303–311.

Clemen, C.S., Marko, M., Strucksberg, K.H., Behrens, J., Wittig, I., Gartner, L., Winter, L., Chevessier, F., Matthias, J., Turk, M., et al. (2015). VCP and PSMF1: Antagonistic regulators of proteasome activity. Biochemical and biophysical research communications 463, 1210–1217.

Coleman, M.P., and Perry, V.H. (2002). Axon pathology in neurological disease: a neglected therapeutic target. Trends in neurosciences 25, 532–537.

Collins, G.A., and Goldberg, A.L. (2017). The Logic of the 26S Proteasome. Cell 169, 792–806.

Conedera, S., Apaydin, H., Li, Y., Yoshino, H., Ikeda, A., Matsushima, T., Funayama, M., Nishioka, K., and Hattori, N. (2016). FBXO7 mutations in Parkinson’s disease and multiple system atrophy. Neurobiology of aging 40, 192 e191–192 e195.

Cox, J., Hein, M.Y., Luber, C.A., Paron, I., Nagaraj, N., and Mann, M. (2014). Accurate proteome-wide label-free quantification by delayed normalization and maximal peptide ratio extraction, termed MaxLFQ. Molecular & cellular proteomics: MCP 13, 2513– 2526.

Cuenda, A., and Rousseau, S. (2007). p38 MAP-kinases pathway regulation, function and role in human diseases. Biochimica et biophysica acta 1773, 1358–1375.

De La Mota-Peynado, A., Lee, S.Y., Pierce, B.M., Wani, P., Singh, C.R., and Roelofs, J. (2013). The proteasome-associated protein Ecm29 inhibits proteasomal ATPase activity and in vivo protein degradation by the proteasome. The Journal of biological chemistry 288, 29467–29481.

de Vrij, F.M., Fischer, D.F., van Leeuwen, F.W., and Hol, E.M. (2004). Protein quality control in Alzheimer’s disease by the ubiquitin proteasome system. Prog Neurobiol 74, 249–270.

Di Fonzo, A., Dekker, M.C., Montagna, P., Baruzzi, A., Yonova, E.H., Correia Guedes, L., Szczerbinska, A., Zhao, T., Dubbel-Hulsman, L.O., Wouters, C.H., et al. (2009). FBXO7 mutations cause autosomal recessive, early-onset parkinsonian-pyramidal syndrome. Neurology 72, 240–245.

DiAntonio, A., Haghighi, A.P., Portman, S.L., Lee, J.D., Amaranto, A.M., and Goodman, C.S. (2001). Ubiquitination-dependent mechanisms regulate synaptic growth and function. Nature 412, 449–452.

Dikic, I., and Elazar, Z. (2018). Mechanism and medical implications of mammalian autophagy. Nature reviews Molecular cell biology 19, 349–364.

Ding, M., Chao, D., Wang, G., and Shen, K. (2007). Spatial regulation of an E3 ubiquitin ligase directs selective synapse elimination. Science 317, 947–951.

Djakovic, S.N., Marquez-Lona, E.M., Jakawich, S.K., Wright, R., Chu, C., Sutton, M.A., and Patrick, G.N. (2012). Phosphorylation of Rpt6 regulates synaptic strength in hippocampal neurons. The Journal of neuroscience: the official journal of the Society for Neuroscience 32, 5126–5131.

Djakovic, S.N., Schwarz, L.A., Barylko, B., DeMartino, G.N., and Patrick, G.N. (2009). Regulation of the proteasome by neuronal activity and calcium/calmodulin-dependent protein kinase II. The Journal of biological chemistry 284, 26655–26665.

Ehlers, M.D. (2003). Activity level controls postsynaptic composition and signaling via the ubiquitin-proteasome system. Nature neuroscience 6, 231–242.

Erturk, A., Wang, Y., and Sheng, M. (2014). Local pruning of dendrites and spines by caspase-3-dependent and proteasome-limited mechanisms. The Journal of neuroscience: the official journal of the Society for Neuroscience 34, 1672–1688.

Finley, D. (2009). Recognition and processing of ubiquitin-protein conjugates by the proteasome. Annual review of biochemistry 78, 477–513.

Finley, D., Chen, X., and Walters, K.J. (2016). Gates, Channels, and Switches: Elements of the Proteasome Machine. Trends in biochemical sciences 41, 77–93.

Gendron, T.F., and Petrucelli, L. (2009). The role of tau in neurodegeneration. Molecular neurodegeneration 4, 13.

Giese, K.P., and Mizuno, K. (2013). The roles of protein kinases in learning and memory. Learning & memory 20, 540–552.

Glickman, M.H., and Ciechanover, A. (2002). The ubiquitin-proteasome proteolytic pathway: destruction for the sake of construction. Physiological reviews 82, 373–428.

Goldberg, A.L. (2003). Protein degradation and protection against misfolded or damaged proteins. Nature 426, 895–899.

Gorbea, C., Pratt, G., Ustrell, V., Bell, R., Sahasrabudhe, S., Hughes, R.E., and Rechsteiner, M. (2010). A protein interaction network for Ecm29 links the 26 S proteasome to molecular motors and endosomal components. The Journal of biological chemistry 285, 31616–31633.

Gratz, S.J., Ukken, F.P., Rubinstein, C.D., Thiede, G., Donohue, L.K., Cummings, A.M., and O’Connor-Giles, K.M. (2014). Highly specific and efficient CRISPR/Cas9-catalyzed homology-directed repair in Drosophila. Genetics 196, 961–971.

Hamilton, A.M., and Zito, K. (2013). Breaking it down: the ubiquitin proteasome system in neuronal morphogenesis. Neural Plast 2013, 196848.

Han, J., Lee, J.D., Bibbs, L., and Ulevitch, R.J. (1994). A MAP kinase targeted by endotoxin and hyperosmolarity in mammalian cells. Science 265, 808–811.

Hegde, A.N., Haynes, K.A., Bach, S.V., and Beckelman, B.C. (2014). Local ubiquitin- proteasome-mediated proteolysis and long-term synaptic plasticity. Frontiers in molecular neuroscience 7, 96.

Heigwer, F., Kerr, G., and Boutros, M. (2014). E-CRISP: fast CRISPR target site identification. Nature methods 11, 122–123.

Hershko, A., and Ciechanover, A. (1998). The ubiquitin system. Annual review of biochemistry 67, 425–479.

Hirokawa, N., Niwa, S., and Tanaka, Y. (2010). Molecular motors in neurons: transport mechanisms and roles in brain function, development, and disease. Neuron 68, 610–638.

Hoover, B.R., Reed, M.N., Su, J., Penrod, R.D., Kotilinek, L.A., Grant, M.K., Pitstick, R., Carlson, G.A., Lanier, L.M., Yuan, L.L., et al. (2010). Tau mislocalization to dendritic spines mediates synaptic dysfunction independently of neurodegeneration. Neuron 68, 1067–1081.

Hsu, M.T., Guo, C.L., Liou, A.Y., Chang, T.Y., Ng, M.C., Florea, B.I., Overkleeft, H.S., Wu, Y.L., Liao, J.C., and Cheng, P.L. (2015). Stage-Dependent Axon Transport of Proteasomes Contributes to Axon Development. Developmental cell 35, 418–431.

Irvine, G.B., El-Agnaf, O.M., Shankar, G.M., and Walsh, D.M. (2008). Protein aggregation in the brain: the molecular basis for Alzheimer’s and Parkinson’s diseases. Mol Med 14, 451–464.

Ittner, L.M., Ke, Y.D., Delerue, F., Bi, M., Gladbach, A., van Eersel, J., Wolfing, H., Chieng, B.C., Christie, M.J., Napier, I.A., et al. (2010). Dendritic function of tau mediates amyloid-beta toxicity in Alzheimer’s disease mouse models. Cell 142, 387–397.

Jan, L.Y., and Jan, Y.N. (1982). Antibodies to horseradish peroxidase as specific neuronal markers in Drosophila and in grasshopper embryos. Proceedings of the National Academy of Sciences of the United States of America 79, 2700–2704.

Jha, S.K., Jha, N.K., Kar, R., Ambasta, R.K., and Kumar, P. (2015). p38 MAPK and PI3K/AKT Signalling Cascades inParkinson’s Disease. International journal of molecular and cellular medicine 4, 67–86.

Johnson, J.O., Mandrioli, J., Benatar, M., Abramzon, Y., Van Deerlin, V.M., Trojanowski, J.Q., Gibbs, J.R., Brunetti, M., Gronka, S., Wuu, J., et al. (2010). Exome sequencing reveals VCP mutations as a cause of familial ALS. Neuron 68, 857–864.

Kall, L., Canterbury, J.D., Weston, J., Noble, W.S., and MacCoss, M.J. (2007). Semisupervised learning for peptide identification from shotgun proteomics datasets. Nature methods 4, 923–925.

Kim, E.K., and Choi, E.J. (2010). Pathological roles of MAPK signaling pathways in human diseases. Biochimica et biophysica acta 1802, 396–405.

Kipreos, E.T., and Pagano, M. (2000). The F-box protein family. Genome biology 1, REVIEWS3002.

Kirk, R., Laman, H., Knowles, P.P., Murray-Rust, J., Lomonosov, M., Mezianeel, K., and McDonald, N.Q. (2008). Structure of a conserved dimerization domain within the Fbox protein Fbxo7 and the PI31 proteasome inhibitor. The Journal of biological chemistry 283, 22325–22335.

Kittel, R.J., Wichmann, C., Rasse, T.M., Fouquet, W., Schmidt, M., Schmid, A., Wagh, D.A., Pawlu, C., Kellner, R.R., Willig, K.I., et al. (2006). Bruchpilot promotes active zone assembly, Ca2+ channel clustering, and vesicle release. Science 312, 1051–1054.

Koppers, M., van Blitterswijk, M.M., Vlam, L., Rowicka, P.A., van Vught, P.W., Groen, E.J., Spliet, W.G., Engelen-Lee, J., Schelhaas, H.J., de Visser, M., et al. (2012). VCP mutations in familial and sporadic amyotrophic lateral sclerosis. Neurobiology of aging 33, 837 e837–813.

Koushik, S.V., Chen, H., Thaler, C., Puhl, H.L., 3rd, and Vogel, S.S. (2006). Cerulean, Venus, and VenusY67C FRET reference standards. Biophysical journal 91, L99–L101.

Kreko-Pierce, T., and Eaton, B.A. (2017). The Drosophila LC8 homolog cut up specifies the axonal transport of proteasomes. Journal of cell science 130, 3388–3398.

Kuo, C.T., Jan, L.Y., and Jan, Y.N. (2005). Dendrite-specific remodeling of Drosophila sensory neurons requires matrix metalloproteases, ubiquitin-proteasome, and ecdysone signaling. Proceedings of the National Academy of Sciences of the United States of America 102, 15230–15235.

Labbadia, J., and Morimoto, R.I. (2015). The biology of proteostasis in aging and disease. Annual review of biochemistry 84, 435–464.

Lecker, S.H., Solomon, V., Mitch, W.E., and Goldberg, A.L. (1999). Muscle protein breakdown and the critical role of the ubiquitin-proteasome pathway in normal and disease states. The Journal of nutrition 129, 227S–237S.

Lee, J.K., and Kim, N.J. (2017). Recent Advances in the Inhibition of p38 MAPK as a Potential Strategy for the Treatment of Alzheimer’s Disease. Molecules 22.

Lee, S.H., Choi, J.H., Lee, N., Lee, H.R., Kim, J.I., Yu, N.K., Choi, S.L., Lee, S.H., Kim, H., and Kaang, B.K. (2008). Synaptic protein degradation underlies destabilization of retrieved fear memory. Science 319, 1253–1256.

Lee, S.Y., De la Mota-Peynado, A., and Roelofs, J. (2011). Loss of Rpt5 protein interactions with the core particle and Nas2 protein causes the formation of faulty proteasomes that are inhibited by Ecm29 protein. The Journal of biological chemistry 286, 36641–36651.

Leggett, D.S., Hanna, J., Borodovsky, A., Crosas, B., Schmidt, M., Baker, R.T., Walz, T., Ploegh, H., and Finley, D. (2002). Multiple associated proteins regulate proteasome structure and function. Molecular cell 10, 495–507.

Lehmann, A., Niewienda, A., Jechow, K., Janek, K., and Enenkel, C. (2010). Ecm29 fulfils quality control functions in proteasome assembly. Molecular cell 38, 879–888.

Levine, B., and Kroemer, G. (2008). Autophagy in the pathogenesis of disease. Cell 132, 27–42.

Li, X., Thompson, D., Kumar, B., and DeMartino, G.N. (2014). Molecular and cellular roles of PI31 (PSMF1) protein in regulation of proteasome function. The Journal of biological chemistry 289, 17392–17405.

Li, X.J., and Li, S. (2011). Proteasomal dysfunction in aging and Huntington disease. Neurobiology of disease 43, 4–8.

Liu, C.W., Li, X., Thompson, D., Wooding, K., Chang, T.L., Tang, Z., Yu, H., Thomas, P.J., and DeMartino, G.N. (2006). ATP binding and ATP hydrolysis play distinct roles in the function of 26S proteasome. Molecular cell 24, 39–50.

Marrus, S.B., Portman, S.L., Allen, M.J., Moffat, K.G., and DiAntonio, A. (2004). Differential localization of glutamate receptor subunits at the Drosophila neuromuscular junction. The Journal of neuroscience: the official journal of the Society for Neuroscience 24, 1406–1415.

McCutchen-Maloney, S.L., Matsuda, K., Shimbara, N., Binns, D.D., Tanaka, K., Slaughter, C.A., and DeMartino, G.N. (2000). cDNA cloning, expression, and functional characterization of PI31, a proline-rich inhibitor of the proteasome. The Journal of biological chemistry 275, 18557–18565.

Mori, H., Kondo, J., and Ihara, Y. (1987). Ubiquitin is a component of paired helical filaments in Alzheimer’s disease. Science 235, 1641–1644.

Morimoto, R.I. (2008). Proteotoxic stress and inducible chaperone networks in neurodegenerative disease and aging. Genes & development 22, 1427–1438.

Morishima-Kawashima, M., Hasegawa, M., Takio, K., Suzuki, M., Titani, K., and Ihara, Y. (1993). Ubiquitin is conjugated with amino-terminally processed tau in paired helical filaments. Neuron 10, 1151–1160.

Munoz, L., and Ammit, A.J. (2010). Targeting p38 MAPK pathway for the treatment of Alzheimer’s disease. Neuropharmacology 58, 561–568.

Murata, S., Yashiroda, H., and Tanaka, K. (2009). Molecular mechanisms of proteasome assembly. Nature reviews Molecular cell biology 10, 104–115.

Nakatogawa, H., Suzuki, K., Kamada, Y., and Ohsumi, Y. (2009). Dynamics and diversity in autophagy mechanisms: lessons from yeast. Nature reviews Molecular cell biology 10, 458–467.

Oddo, S. (2008). The ubiquitin-proteasome system in Alzheimer’s disease. J Cell Mol Med 12, 363–373.

Olenych, S.G., Claxton, N.S., Ottenberg, G.K., and Davidson, M.W. (2007). The fluorescent protein color palette. Current protocols in cell biology Chapter 21, Unit 21 25.

Otero, M.G., Alloatti, M., Cromberg, L.E., Almenar-Queralt, A., Encalada, S.E., Pozo Devoto, V.M., Bruno, L., Goldstein, L.S., and Falzone, T.L. (2014). Fast axonal transport of the proteasome complex depends on membrane interaction and molecular motor function. Journal of cell science 127, 1537–1549.

Owald, D., and Sigrist, S.J. (2009). Assembling the presynaptic active zone. Current opinion in neurobiology 19, 311–318.

Paisan-Ruiz, C., Guevara, R., Federoff, M., Hanagasi, H., Sina, F., Elahi, E., Schneider, S.A., Schwingenschuh, P., Bajaj, N., Emre, M., et al. (2010). Early-onset L-dopa-responsive parkinsonism with pyramidal signs due to ATP13A2, PLA2G6, FBXO7 and spatacsin mutations. Movement disorders: official journal of the Movement Disorder Society 25, 1791–1800.

Pak, D.T., and Sheng, M. (2003). Targeted protein degradation and synapse remodeling by an inducible protein kinase. Science 302, 1368–1373.

Park, S., Kim, W., Tian, G., Gygi, S.P., and Finley, D. (2011). Structural defects in the regulatory particle-core particle interface of the proteasome induce a novel proteasome stress response. The Journal of biological chemistry 286, 36652–36666.

Patrick, G.N. (2006). Synapse formation and plasticity: recent insights from the perspective of the ubiquitin proteasome system. Current opinion in neurobiology 16, 90– 94.

Perry, G., Friedman, R., Shaw, G., and Chau, V. (1987). Ubiquitin is detected in neurofibrillary tangles and senile plaque neurites of Alzheimer disease brains. Proceedings of the National Academy of Sciences of the United States of America 84, 3033–3036.

Pfeiffer, B.D., Jenett, A., Hammonds, A.S., Ngo, T.T., Misra, S., Murphy, C., Scully, A., Carlson, J.W., Wan, K.H., Laverty, T.R., et al. (2008). Tools for neuroanatomy and neurogenetics in Drosophila. Proceedings of the National Academy of Sciences of the United States of America 105, 9715–9720.

Pfeiffer, B.D., Ngo, T.T., Hibbard, K.L., Murphy, C., Jenett, A., Truman, J.W., and Rubin, G.M. (2010). Refinement of tools for targeted gene expression in Drosophila. Genetics 186, 735–755.

Port, F., Chen, H.M., Lee, T., and Bullock, S.L. (2014). Optimized CRISPR/Cas tools for efficient germline and somatic genome engineering in Drosophila. Proceedings of the National Academy of Sciences of the United States of America 111, E2967–2976.

Qian, M.X., Pang, Y., Liu, C.H., Haratake, K., Du, B.Y., Ji, D.Y., Wang, G.F., Zhu, Q.Q., Song, W., Yu, Y., et al. (2013). Acetylation-mediated proteasomal degradation of core histones during DNA repair and spermatogenesis. Cell 153, 1012–1024.

Ramachandran, K.V., Fu, J.M., Schaffer, T.B., Na, C.H., Delannoy, M., and Margolis, S.S. (2018). Activity-Dependent Degradation of the Nascentome by the Neuronal Membrane Proteasome. Molecular cell 71, 169–177 e166.

Ramachandran, K.V., and Margolis, S.S. (2017). A mammalian nervous-system-specific plasma membrane proteasome complex that modulates neuronal function. Nature structural & molecular biology 24, 419–430.

Rappsilber, J., Mann, M., and Ishihama, Y. (2007). Protocol for micro-purification, enrichment, pre-fractionation and storage of peptides for proteomics using StageTips. Nature protocols 2, 1896–1906.

Reck-Peterson, S.L., Redwine, W.B., Vale, R.D., and Carter, A.P. (2018). The cytoplasmic dynein transport machinery and its many cargoes. Nature reviews Molecular cell biology 19, 382–398.

Ren, X., Sun, J., Housden, B.E., Hu, Y., Roesel, C., Lin, S., Liu, L.P., Yang, Z., Mao, D., Sun, L., et al. (2013). Optimized gene editing technology for Drosophila melanogaster using germ line-specific Cas9. Proceedings of the National Academy of Sciences of the United States of America 110, 19012–19017.

Ross, C.A., and Poirier, M.A. (2004). Protein aggregation and neurodegenerative disease. Nature medicine 10 Suppl, S10–17.

Rubinsztein, D.C. (2006). The roles of intracellular protein-degradation pathways in neurodegeneration. Nature 443, 780–786.

Sandri, M. (2013). Protein breakdown in muscle wasting: role of autophagy-lysosome and ubiquitin-proteasome. The international journal of biochemistry & cell biology 45, 2121–2129.

Schindelin, J., Arganda-Carreras, I., Frise, E., Kaynig, V., Longair, M., Pietzsch, T., Preibisch, S., Rueden, C., Saalfeld, S., Schmid, B., et al. (2012). Fiji: an open-source platform for biological-image analysis. Nature methods 9, 676–682.

Schliwa, M., and Woehlke, G. (2003). Molecular motors. Nature 422, 759–765.

Schmidt, M., and Finley, D. (2014). Regulation of proteasome activity in health and disease. Biochimica et biophysica acta 1843, 13–25.

Schwanhausser, B., Busse, D., Li, N., Dittmar, G., Schuchhardt, J., Wolf, J., Chen, W., and Selbach, M. (2011). Global quantification of mammalian gene expression control. Nature 473, 337–342.

Sherva, R., Baldwin, C.T., Inzelberg, R., Vardarajan, B., Cupples, L.A., Lunetta, K., Bowirrat, A., Naj, A., Pericak-Vance, M., Friedland, R.P., et al. (2011). Identification of novel candidate genes for Alzheimer’s disease by autozygosity mapping using genome wide SNP data. Journal of Alzheimer’s disease: JAD 23, 349–359.

Skaar, J.R., Pagan, J.K., and Pagano, M. (2013). Mechanisms and function of substrate recruitment by F-box proteins. Nat Rev Mol Cell Biol 14, 369–381.

Skowyra, D., Craig, K.L., Tyers, M., Elledge, S.J., and Harper, J.W. (1997). F-box proteins are receptors that recruit phosphorylated substrates to the SCF ubiquitin-ligase complex. Cell 91, 209–219.

Speese, S.D., Trotta, N., Rodesch, C.K., Aravamudan, B., and Broadie, K. (2003). The ubiquitin proteasome system acutely regulates presynaptic protein turnover and synaptic efficacy. Current biology: CB 13, 899–910.

Sun, B., Xu, P., and Salvaterra, P.M. (1999). Dynamic visualization of nervous system in live Drosophila. Proceedings of the National Academy of Sciences of the United States of America 96, 10438–10443.

Tai, H.C., and Schuman, E.M. (2008). Ubiquitin, the proteasome and protein degradation in neuronal function and dysfunction. Nature reviews Neuroscience 9, 826–838.

Taus, T., Kocher, T., Pichler, P., Paschke, C., Schmidt, A., Henrich, C., and Mechtler, K. (2011). Universal and confident phosphorylation site localization using phosphoRS. Journal of proteome research 10, 5354–5362.

Tyanova, S., Temu, T., Sinitcyn, P., Carlson, A., Hein, M.Y., Geiger, T., Mann, M., and Cox, J. (2016). The Perseus computational platform for comprehensive analysis of (prote)omics data. Nature methods 13, 731–740.

Valakh, V., Naylor, S.A., Berns, D.S., and DiAntonio, A. (2012). A large-scale RNAi screen identifies functional classes of genes shaping synaptic development and maintenance. Developmental biology 366, 163–171.

Vale, R.D. (2003). The molecular motor toolbox for intracellular transport. Cell 112, 467– 480.

Varshavsky, A. (2005). Regulated protein degradation. Trends in biochemical sciences 30, 283–286.

Varshavsky, A. (2012). The ubiquitin system, an immense realm. Annual review of biochemistry 81, 167–176.

Verdoes, M., Florea, B.I., Menendez-Benito, V., Maynard, C.J., Witte, M.D., van der Linden, W.A., van den Nieuwendijk, A.M., Hofmann, T., Berkers, C.R., van Leeuwen, F.W., et al. (2006). A fluorescent broad-spectrum proteasome inhibitor for labeling proteasomes in vitro and in vivo. Chemistry & biology 13, 1217–1226.

Vilchez, D., Saez, I., and Dillin, A. (2014). The role of protein clearance mechanisms in organismal ageing and age-related diseases. Nature communications 5, 5659.

Vingill, S., Brockelt, D., Lancelin, C., Tatenhorst, L., Dontcheva, G., Preisinger, C., Schwedhelm-Domeyer, N., Joseph, S., Mitkovski, M., Goebbels, S., et al. (2016). Loss of FBXO7 (PARK15) results in reduced proteasome activity and models a parkinsonism-like phenotype in mice. The EMBO journal 35, 2008–2025.

Wagh, D.A., Rasse, T.M., Asan, E., Hofbauer, A., Schwenkert, I., Durrbeck, H., Buchner, S., Dabauvalle, M.C., Schmidt, M., Qin, G., et al. (2006). Bruchpilot, a protein with homology to ELKS/CAST, is required for structural integrity and function of synaptic active zones in Drosophila. Neuron 49, 833–844.

Wagih, O., Sugiyama, N., Ishihama, Y., and Beltrao, P. (2016). Uncovering Phosphorylation-Based Specificities through Functional Interaction Networks. Molecular & cellular proteomics: MCP 15, 236–245.

Wan, H.I., DiAntonio, A., Fetter, R.D., Bergstrom, K., Strauss, R., and Goodman, C.S. (2000). Highwire regulates synaptic growth in Drosophila. Neuron 26, 313–329.

Wang, X., Yen, J., Kaiser, P., and Huang, L. (2010). Regulation of the 26S proteasome complex during oxidative stress. Science signaling 3, ra88.

Watts, R.J., Hoopfer, E.D., and Luo, L. (2003). Axon pruning during Drosophila metamorphosis: evidence for local degeneration and requirement of the ubiquitin-proteasome system. Neuron 38, 871–885.

Willeumier, K., Pulst, S.M., and Schweizer, F.E. (2006). Proteasome inhibition triggers activity-dependent increase in the size of the recycling vesicle pool in cultured hippocampal neurons. The Journal of neuroscience: the official journal of the Society for Neuroscience 26, 11333–11341.

Wolff, S., Weissman, J.S., and Dillin, A. (2014). Differential scales of protein quality control. Cell 157, 52–64.

Wu, J.S., and Luo, L. (2006). A protocol for mosaic analysis with a repressible cell marker (MARCM) in Drosophila. Nature protocols 1, 2583–2589.

Yashiroda, H., Toda, Y., Otsu, S., Takagi, K., Mizushima, T., and Murata, S. (2015). N-terminal alpha7 deletion of the proteasome 20S core particle substitutes for yeast PI31 function. Molecular and cellular biology 35, 141–152.

Yasuda, S., Sugiura, H., Tanaka, H., Takigami, S., and Yamagata, K. (2011). p38 MAP kinase inhibitors as potential therapeutic drugs for neural diseases. Central nervous system agents in medicinal chemistry 11, 45–59.

Yi, J.J., and Ehlers, M.D. (2005). Ubiquitin and protein turnover in synapse function. Neuron 47, 629–632.

Yoshiyama, Y., Higuchi, M., Zhang, B., Huang, S.M., Iwata, N., Saido, T.C., Maeda, J., Suhara, T., Trojanowski, J.Q., and Lee, V.M. (2007). Synapse loss and microglial activation precede tangles in a P301S tauopathy mouse model. Neuron 53, 337–351.

Yu, W., and Lu, B. (2012). Synapses and dendritic spines as pathogenic targets in Alzheimer’s disease. Neural Plast 2012, 247150.

Zaiss, D.M., Standera, S., Holzhutter, H., Kloetzel, P., and Sijts, A.J. (1999). The proteasome inhibitor PI31 competes with PA28 for binding to 20S proteasomes. FEBS letters 457, 333–338.

Zarubin, T., and Han, J. (2005). Activation and signaling of the p38 MAP kinase pathway. Cell research 15, 11–18.

Zhao, Y., Hegde, A.N., and Martin, K.C. (2003). The ubiquitin proteasome system functions as an inhibitory constraint on synaptic strengthening. Current biology: CB 13, 887–898.

Zhong, L., and Belote, J.M. (2007). The testis-specific proteasome subunit Prosalpha6T of D. melanogaster is required for individualization and nuclear maturation during spermatogenesis. Development 134, 3517–3525.

Zhu, G., Fujii, K., Belkina, N., Liu, Y., James, M., Herrero, J., and Shaw, S. (2005). Exceptional disfavor for proline at the P + 1 position among AGC and CAMK kinases establishes reciprocal specificity between them and the proline-directed kinases. The Journal of biological chemistry 280, 10743–10748.

Zoghbi, H.Y., and Orr, H.T. (2000). Glutamine repeats and neurodegeneration. Annual review of neuroscience 23, 217–247.

